# Improved precision, sensitivity, and adaptability of Ordered Two-Template Relay cDNA library preparation for RNA sequencing

**DOI:** 10.1101/2024.11.09.622813

**Authors:** Lucas Ferguson, Heather E. Upton, Sydney C. Pimentel, Chris Jeans, Nicholas T. Ingolia, Kathleen Collins

**Affiliations:** Department of Molecular and Cell Biology, University of California, Berkeley, Berkeley, USA; Center for Computational Biology, University of California, Berkeley, Berkeley, CA, USA; MacroLab, University of California, Berkeley, Berkeley, CA, USA; California Institute for Quantitative Biosciences, University of California, Berkeley, Berkeley, USA; Addition Therapeutics, 201 Haskins Way, South San Francisco, CA 94080; NYU Grossman School of Medicine, 550 First Avenue, New York, NY 10016

**Keywords:** non-coding RNA, reverse transcriptase, template jumping, non-templated nucleotide addition, terminal transferase, OTTR

## Abstract

Sequencing RNAs that are biologically processed or degraded to less than ∼100 nucleotides typically involves multi-step, low-yield protocols with bias and information loss inherent to ligation and/or polynucleotide tailing. We recently introduced Ordered Two-Template Relay (OTTR), a method that captures obligatorily end-to-end sequences of input molecules and, in the same reverse transcription step, also appends 5′ and 3′ sequencing adapters of choice. OTTR has been thoroughly benchmarked for optimal production of microRNA, tRNA and tRNA fragments, and ribosome-protected mRNA footprint libraries. Here we sought to characterize, quantify, and ameliorate any remaining bias or imprecision in the end-to-end capture of RNA sequences. We introduce new metrics for the evaluation of sequence capture and use them to optimize reaction buffers, reverse transcriptase sequence, adapter oligonucleotides, and overall workflow. Modifications of the reverse transcriptase and adapter oligonucleotides increased the 3′ and 5′ end-precision of sequence capture and minimized overall library bias. Improvements in recombinant expression and purification of the truncated *Bombyx mori* R2 reverse transcriptase used in OTTR reduced non-productive sequencing reads by minimizing bacterial nucleic acids that compete with low-input RNA molecules for cDNA synthesis, such that with miRNA input of 3 picograms (less than 1 fmol), fewer than 10% of sequencing reads are bacterial nucleic acid contaminants. We also introduce a rapid, automation-compatible OTTR protocol that enables gel-free, length-agnostic enrichment of cDNA duplexes from unwanted adapter-only side products. Overall, this work informs considerations for unbiased end-to-end capture and annotation of RNAs independent of their sequence, structure, or post-transcriptional modifications.

## Introduction

Nearly two decades of innovations in polyadenylated messenger (m) RNA sequencing have generated numerous methods for the bulk and single-cell sequencing of protein-coding transcripts (Vandereyken et al. 2023). In comparison, methods that capture the full complexity of non-coding RNAs (ncRNA) or processed small RNAs remain underdeveloped (Tosar et al. 2024) and comprehensively profiling these RNAs at low input, for example single-cell profiling, is an unmet challenge (Wang et al. 2019, Hücker et al. 2021). Although mRNA sequencing is an unquestionably useful tool for assessing cell state, profiles of microRNAs (miRNA), transfer RNAs (tRNA), tRNA-derived fragments (tRF), vesicle-trafficked and lysosomal RNA fragments, and non-polyadenylated pathogen-derived RNAs could be additionally or even more informative (Muthukumar et al. 2024, Shi et al. 2022, O’Brien et al. 2020). Increasingly, ncRNAs are recognized as important pre- and post-transcriptional regulators of gene expression, with emerging roles in epigenetic inheritance and cell-to-cell communication (Kaikkonen et al. 2011, Boskovic et al. 2020, van Niel et al. 2022). Without widespread adoption of sensitive, accurate, and reproducible methods to inventory these transcripts, the quantitative appraisal of biology is incomplete.

To fill this knowledge gap, complementary (c) DNA library preparation methods that do not require specific RNA sequence motifs and are not impaired by RNA structure or modified ribonucleotides are needed. Retroviral reverse transcriptases (RTs) exclusively initiate cDNA synthesis by annealing a cognate primer to a sequence in the RNA template, *e.g.,* in polydeoxythymidine-primed cDNA synthesis from mRNA. To make cDNA libraries from small RNAs, which do not possess a shared 3′ sequence, the 3′ ends of RNAs are typically first either tailed with adenosines by enzymes such as *Escherichia coli* or yeast polyadenosine polymerase (yPAP), or ligated to an RNA adapter using specialized ligase enzymes (Lu et al. 2007, Coenen-Stass et al. 2018, Androvic et al 2022). Both 3′-standardization methods can be inefficient and favor the capture of specific RNAs based on sequence or structure (Yehudai-Resheff and Schuster 2000, Fuchs et al. 2015, Tunney et al. 2018, Ferguson et al. 2023). An approach to extend cDNA with an adapter sequence is to rely on the RT to perform multiple non-templated nucleotide additions (NTA) to the cDNA 3′ end. As first developed for cDNA library synthesis using the retroviral Moloney murine leukemia virus RT, these NTA nucleotides can support base pairing with the 3′ end of an oligonucleotide adapter template and thereby prime continued cDNA synthesis (Wulf et al. 2019, Zhu et al. 2001).

A similar but distinct “template jumping” activity has been observed using non-retroviral RTs, which can initiate cDNA synthesis from a blunt-ended primer duplex or one with a single-nucleotide 3′ overhang. This is exploited in thermostable group II intron RT cDNA library synthesis, which uses a DNA/RNA primer duplex with a single, 3′-overhang, degenerate deoxyribonucleotide (dN, *i.e.*, adenosine (A), thymidine (T), guanosine (G), or cytidine (C)) to capture RNAs without a standardized 3′ sequence (Xu et al. 2019). Recently we exploited the particularly robust template-jumping activity of the non-long terminal repeat (non-LTR) retrotransposon protein encoded by *Bombyx mori* R2 (Bibiłło and Eickbush 2002a, Bibiłło and Eickbush 2002b) to establish a new cDNA library synthesis method (Upton et al. 2021). The *B. mori* R2 RT can initiate reverse transcription on RNA or DNA templates using a duplexed primer with 3′ end that is blunt or has a +1 or at most a +2 3′-overhang (hereafter +1 or +2). *B. mori* R2 RT, and enzymes engineered from it, readily synthesize cDNA concatemers from multiple physically separate input templates molecules via back-to-back template jumps (Bibiłło et al. 2002a, Upton et al. 2021, Pimentel et al. 2022). The capacity to initiate reverse transcription in this way may have evolved with the non-LTR retrotransposon insertion mechanism of target-primed reverse transcription—in this process the retrotransposon-protein endonuclease domain nicks the target site, liberating a 3′-hydroxyl (3′-OH) primer that the RT domain uses to initiate cDNA polymerization at the 3′ end of its bound RNA (Eickbush and Eickbush 2015). Non-LTR retrotransposon RTs have also evolved efficient strand-displacement activity to confer the high processivity needed to successfully copy a long and structured retrotransposon RNA template (Bibiłło et al. 2002b, Kurzynska-Kokorniak et al. 2007).

Using a truncated version of *B. mori* R2 protein, we established Ordered Two-Template Relay (OTTR) to reverse transcribe a continuous cDNA from discontinuous templates. Two types of templates are reverse transcribed in a specific order that ensures each input-template cDNA is flanked on both sides by the Illumina TruSeq sequencing adapters, Read1 and Read2 (Upton et al. 2021). OTTR exploits the manganese-stimulated terminal-transferase (hereafter 3′-labeling) activity of a truncated, endonuclease-inactivated *B. mori* R2 protein (hereafter BoMoC) and a dideoxynucleotide triphosphate (ddNTP) substrate to extend the 3′ ends of single-stranded or double-stranded DNAs, RNAs, or DNA/RNA duplexes (Upton et al. 2021). This 3′-labeling activity differs from NTA in that it is robust only in reactions with non-physiologically high manganese concentrations and the optimal substrate is single-stranded rather than the cDNA duplex extended by NTA (Upton et al. 2021). While other polymerases have also demonstrated manganese-enhanced 3′-labeling activity (Pelletier et al. 1996, Kent et al. 2016, Park et al. 2022, Ohtsubo et al. 2017, Balint and Unk 2024), the divalent-ion-dependent toggle of R2 protein activity from exclusively 3′ labeling in manganese to exclusively templated cDNA synthesis in magnesium is particularly enabling (Upton et al. 2021).

In the first step of OTTR, BoMoC is used in 3′-labeling conditions to append a single nucleotide, either ddA or ddG, collectively dideoxypurine (ddR, *i.e.*, R is either A or G), to the 3′ ends of input nucleic acids (**Fig. 1**, step 1; note that previous OTTR conditions are in black or gray text, and improvements described below are given in red text and mark-out of black text with a red line). Unincorporated ddNTPs are inactivated by recombinant shrimp alkaline phosphatase (rSAP), with subsequent chelation of divalent metal ions by addition of glycol-bis(β-aminoethyl ether)-N,N,N′,N′-tetraacetic acid (EGTA) to minimize RNA hydrolysis during rSAP heat-inactivation (**Fig. 1**, step 2). Then magnesium is added in excess of EGTA (EGTA preferentially chelates manganese and the zinc from rSAP, sparing magnesium) and BoMoC-mediated cDNA reverse transcription is primed by the formation of a single base pair between the 3′ ends of +1 deoxypyrimidine (Y, *i.e.,* Y is either T or C) DNA/RNA primer duplexes and the 3′ ends of 3′ddR-labeled templates (**Fig. 1**, step 3 1^st^ jump). Following cDNA synthesis across the 3′ddR-labeled templates, BoMoC will extend the cDNA/RNA duplex with a +1 G NTA, providing the specificity needed for the second template jump to a 3′ C oligonucleotide that encodes the cDNA 3′ adapter (hereafter adapter template) rather than another RNA from the input pool (**Fig. 1**, step 3 2^nd^ jump). These template jumps convert the input pool into end-to-end transcribed cDNA with flanking 5′ and 3′ adapters. Bulky dye molecules are present on the 5′ end of the primer strand and the adapter template to enable cDNA product detection and to inhibit template jumping from the adapter template 5′ end, respectively.

**Figure 1:**
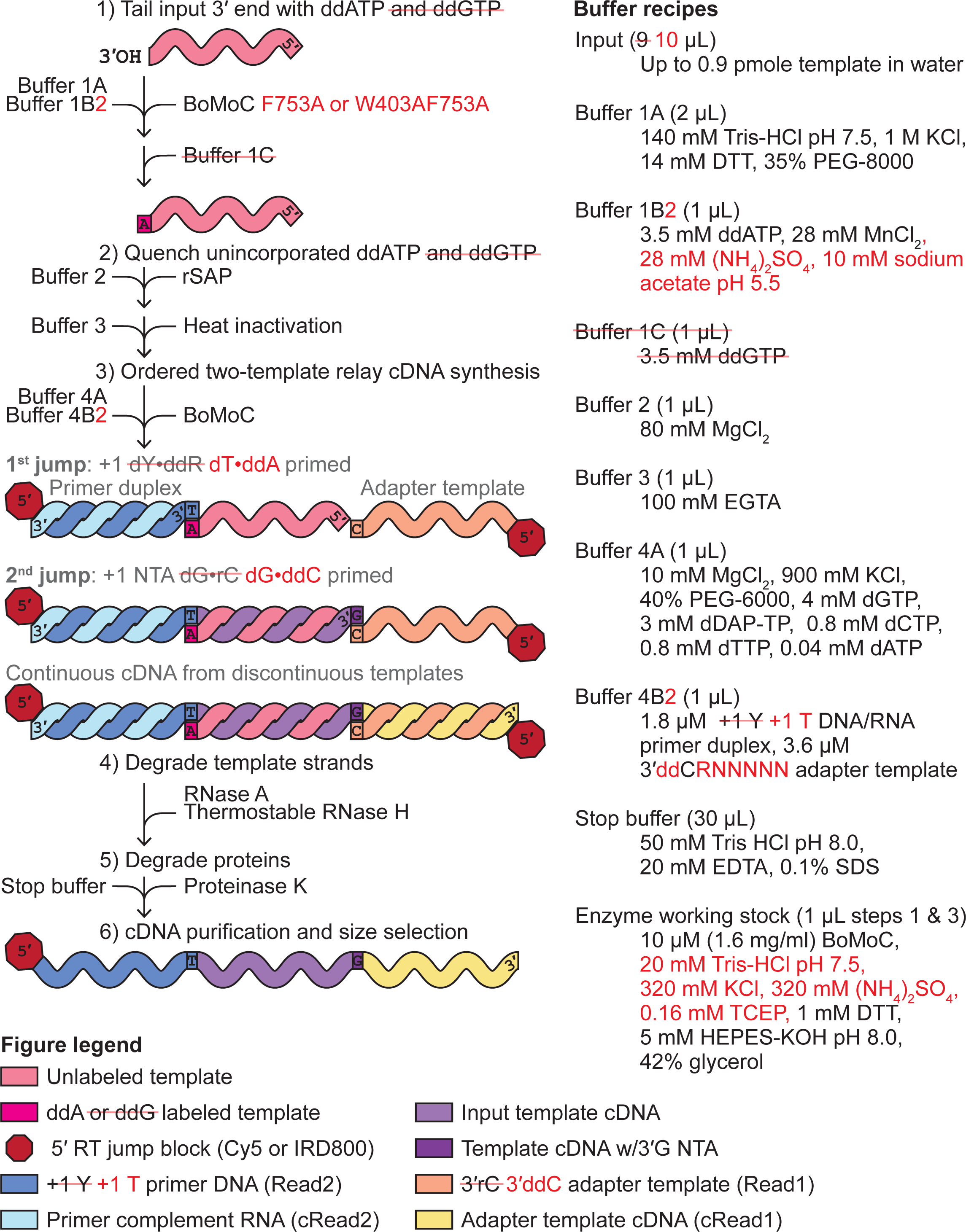
Improvements in the OTTR library synthesis workflow. Previous OTTR conditions are in black or gray text, and improvements are given in red text and mark-out of black text with a red line. The left side explains the steps, while the right side gives buffer recipes.

To date, we have established OTTR as a low-bias cDNA library synthesis method for the high-throughput profiling of miRNA, extracellular RNA, and ribosome-protected mRNA footprint (RPF) sequences (Upton et al. 2021, Ferguson et al. 2023), while others have demonstrated the utility of OTTR for detecting and quantifying the relative abundance of tRNAs and tRFs and their post-transcriptional modifications (Gustafsson et al. 2022, d’Almeida et al. 2023, Manning et al. 2024, Davey-Young et al. 2024). However, we have yet to establish metrics to assess precision of end-to-end sequence capture (*i.e.,* how frequently sequence-capture truncates or adds a terminal nucleotide(s)), determine enzyme and buffer stability with storage, query the lower-input limit for productive library synthesis, include a unique molecular identifier (UMI) to uniquely tag each cDNA molecule, or explore the use of improvements to BoMoC activities via mutagenesis (Pimentel et al. 2022). In this study, we sought to identify sources of bias or information loss in the OTTR protocol and introduce solutions to mitigate these and other concerns.

## Results and Discussion

### Precision of sequence capture at input RNA 3′ ends

In the most recently published version of OTTR (Ferguson et al. 2023; OTTR v1.2), built upon the initial OTTR protocol (Upton et al. 2021; OTTR v1.1), an input RNA 3′-OH is first extended by BoMoC in the 3′-labeling step to gain a 3′ddR, with initial 3′-labeling using ddATP followed by ddGTP chase, before primer duplexes with cognate, single-nucleotide 3′Y overhang initiate reverse transcription (**Fig. 1**). To quantify the extent to which the 3′-labeling step determines input RNA conversion to cDNA, we performed OTTR either without the 3′-labeling step or using yPAP and ddATP for 3′ labeling (Martin and Keller 1998) instead of BoMoC. To assess library bias in OTTR, we made cDNA libraries from the miRXplore equimolar pool of 962 synthetic miRNAs, a reference standard commonly used to benchmark small RNA library synthesis protocols (Coenen-Stass et al. 2018, Shore et al. 2019, Xu et al. 2019, Giraldez et al. 2018, Herbert et al. 2020, Upton et al. 2021). Deviation from equimolar miRNA representation in the sequencing reads was quantified by the library-wide coefficient of variation (CV), a measure of the variation in number of sequencing reads per miRNA from the expected equimolar representation. As a metric for 3′ precision, we scored the fraction of reads for each miRNA that reflected 3′ priming on the added 3′ddR nucleotide instead of an alternative mechanism (**Fig. 2A**). The significance of the ddR 3′-labeling step was evident in the dramatic reduction of both library CV and 3′ precision (mean fraction of 3′-precise alignments across all miRNAs) in the cDNA libraries made without the ddR 3′-labeling step (“Unlabeled"), or when yPAP was used in place of BoMoC (**Fig. 2B**). As expected, this bias was most pronounced when comparing miRNAs with the 3′ ribonucleotide adenosine (rA) or guanosine (rG), collectively rR (*i.e.*, either rA or rG), which were over-represented, to miRNAs with a 3′ uridine (rU) or cytidine (rC), collectively rY (*i.e.*, either rU or rC), which were under-represented (**Supplemental Fig. 1A-B**).

**Figure 2:**
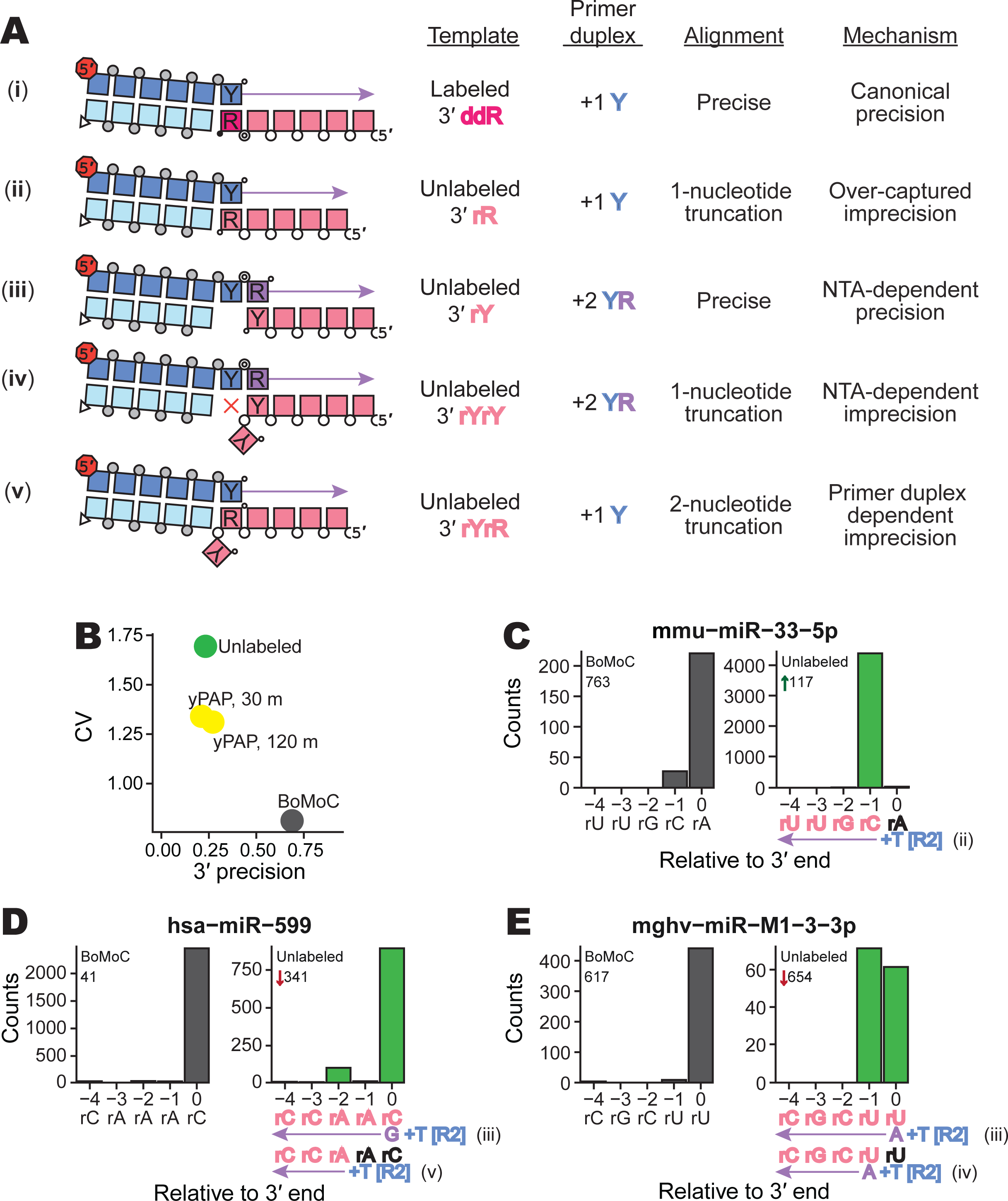
Evaluating imprecision of end-to-end sequence capture at RNA 3′ ends. **A**, Mechanisms, numbered by Roman numeral, of 3′-end capture in OTTR. Column one illustrates five specificities for capturing an input-template 3′ end, following the color legend from Figure 1. Phosphodiester bonds are represented by circles: gray for chemically synthesized bonds, concentric double circle for BoMoC-polymerized, unfilled for input template RNA. Strand 3′ end symbols are a small black circle for dideoxynucleotide, a small open circle for 3′-OH, and a triangle for 3′-OH replacement with an unextendible carbon linker. The red octagon indicates the presence of a 5′ fluorescent dye. Column two specifies the labeling status of the input-template 3′ end. Column three identifies the primer-duplex 3′ overhang supporting a template jump. Column four details the impact on 3′ sequence coverage. Column five categorizes the 3′-end capture mechanism. **B**, Library-wide CV of miRXplore miRNA counts and mean fraction of alignments with complete 3′-end coverage (3′ precision). BoMoC, yPAP (for 30 or 120 minutes), or no enzyme (Unlabeled) was used for 3′ labeling. **C-E**, Panel pairs depict 3′-end coverage with BoMoC (left) or no enzyme (right) for 3′ labeling. The miRNA sequence and name, and relative rank-order, are indicated at top. At bottom, position “0” is the miRNA 3′ end. Right panels have at bottom the miRNA sequence color-coded by alignment inclusion (pink) or exclusion (black). The Read2 (R2) primer sequence and adjacent +1 T overhang (+T) are in blue, and reference to the capture mechanism detailed in (**A**) is indicated. Input RNA 3′ sequence excluded by adapter trimming is indicated in black. The miRNA mmu-miR-33-5p is miRBase: MIMAT0000667, the miRNA hsa-miR-599 is miRBase: MIMAT0003267, and the miRNA mghv-miR-M1-3-3p is miRBase: MIMAT0001566.

Given the poor prognosis on library capture bias when input 3′ labeling was omitted, we were surprised that all but one of the miRNA sequences were captured, with only 27 miRNAs under-represented 100-fold or more from equimolar capture. To investigate how miRNAs were captured into an OTTR cDNA library without 3′ labeling, we compared rank-order of miRNA read abundance relative to 3′ precision. Three types of non-canonical 3′-end capture were detected that resulted in a miRNA 3′ truncation, and we inferred a fourth type indistinguishable from precise 3′-end capture (**Fig. 2A**). First, miRNAs with a 3′rR do not need 3′ labeling to be captured. Input miRNAs with 3′rR that failed to gain a 3′ddR prior to cDNA synthesis have their 3′ ribonucleotide removed by sequence trimming and appear 1 nucleotide shorter in length (**Fig. 2A, ii**). These 3′rR miRNAs became over-represented when 3′ labeling was excluded, with a corresponding under-representation of 3′rY miRNA (**Supplemental Fig. 1A-B**). For example, when 3′ labeling was excluded, mmu-miR-33-5p changed in rank order from 763^rd^ to 117^th^, with the trimmed reads appearing as if the miRNA ended at its penultimate rC rather than 3′rA (**Fig. 2C**). We designated these events as imprecision from “over-capture” (**Fig. 2A**, imprecision mechanism **ii**).

Additional classes of non-canonical 3′-end capture were inferred from analysis of miRNAs with 3′rY that showed apparently precise capture despite omission of a 3′-labeling step (**Supplemental Fig. 1B**). In these cases, 3′rY capture was facilitated due to additional NTA to the +1 Y primer duplex. Because NTA by BoMoC strongly favors use of dR nucleotides (Upton et al. 2021), NTA to the primer duplex thus forms a +2 YR overhang (**Fig. 2A**, **iii** and **iv**), which BoMoC can use to initiate a template jump (Upton et al. 2021, Pimentel et al. 2022). This can account for why miRNAs hsa-miR-599 (**Fig. 2D**) and mghv-miR-M1-3-3p (**Fig. 2E**) appeared to be precisely captured when 3′ labeling was excluded. In addition to the apparently precise capture by use of +2 YR primer (**Fig. 2A**, imprecision mechanism **iii**), for some miRNAs there was imprecise capture from +2 YR primer corresponding to cDNA synthesis initiation on the penultimate ribonucleotide (-1 from the 3′ end, see **Fig. 2E**; product by **Fig. 2A** imprecision mechanism **iv**). The penultimate rA also appeared to be captured, albeit inefficiently, by primer duplexes with the +1 T (**Fig. 2A**, imprecision mechanism **v**, resulting in an apparent two-nucleotide truncation (**Fig. 2D**). Previous template-jumping assays with full-length *B. mori* R2 protein detected cDNA synthesis initiation slightly internal to a template 3′ end when precise 3′-end capture was disfavored by skew of deoxynucleotide triphosphate (dNTP) concentrations (Bibiłło and Eickbush 2004).

We concluded that BoMoC will capture input RNA sequences with imprecision if the 3′ddR is not present, resulting in detection of more RNA sequences than we anticipated for use of the +1 Y primer duplex (**Supplemental Fig. 1A-B**). Since RNAs with 3′rR can be captured efficiently without a 3′ddR label, their relative abundances compared to 3′rY RNAs are inflated if the 3′-labeling step is inefficient, creating a 3′-end-dependent bias of template capture. Moreover, the misappropriation of the +1 Y primer duplex to capture unlabeled RNAs increased the number of alignments with an apparent one-nucleotide, or greater, 3′ truncation due to adapter trimming, *i.e.,* a 3′ imprecision. While this 3′ bias and imprecision can be detected if known sequences are used as input templates, we reasoned that optimization of the 3′-labeling step of OTTR would reduce sequence bias and improve the interpretation of end-to-end sequence of input RNAs that lack a reference standard.

### Input RNA 3′ labeling for optimal 3′ precision

Previous structure/function studies demonstrated that BoMoC side-chain substitutions can increase manganese-stimulated 3′ tailing of single-stranded RNA (Pimentel et al. 2022). Here we compared previously generated BoMoC variants with side-chain substitutions W403A, G415A, F753A, or I770A (Pimentel et al. 2022), and a previously unreported double-mutant W403AF753A, for 3′ labeling of input RNA in the first step of OTTR. OTTR cDNA libraries made by BoMoC W403A, I770A, F753A or W403AF753A had significantly higher 3′ precision (one-sided student’s *t*-test p-value of 2.69 x 10^-2^, 4.06 x 10^-4^, 4.37 x 10^-11^, and 5.52 x 10^-21^, respectively) and lower CV compared to cDNA libraries made using original BoMoC (BoMoC WT) for the 3′-labeling step (**Fig. 3A**). Libraries made using BoMoC F753A or I770A for the 3′-labeling step had the lowest library-wide CV, and libraries made with BoMoC F753A or W403AF753A had the highest 3′ precision (**Fig. 3A**). Hereafter, unless specified otherwise, we used BoMoC F753A or W403AF753A for the 3′-labeling step (**Fig. 1**, changes in red). BoMoC F753A has been previously used for 3′ labeling in ribosome-profiling library synthesis by OTTR (Ferguson et al. 2023, Li et al. 2022, Mestre-Fos et al. 2024).

**Figure 3:**
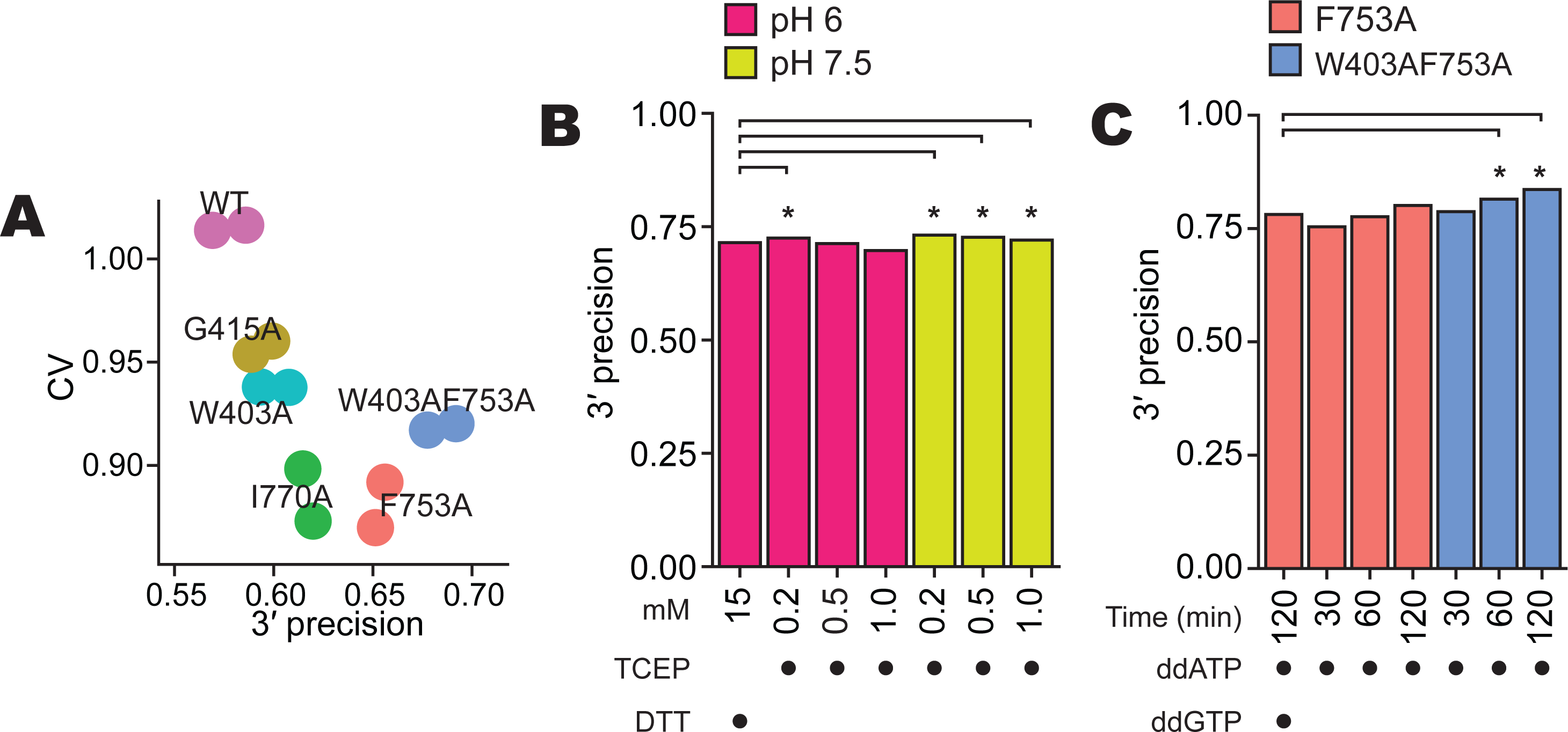
Optimization of 3′ labeling via BoMoC sequence and reaction buffers. **A**, Library-wide CV and 3′ precision of miRXplore miRNA as in Figure 2B, using BoMoC WT or variant enzymes. Replicate cDNA libraries have the same color. **B**, Comparison of 3′ precision under different BoMoC working-stock enzyme storage buffer conditions. Significant differences in 3′ precision (denoted by asterisks) were determined by a paired, two-sided student’s *t*-test for each miRNA, with p-values adjusted by Bonferroni correction. Each library was benchmarked against a library prepared with enzyme diluted in 15 mM DTT in pH 6.0 diluent buffer. **C,** Comparison of 3′ precision using different BoMoC proteins, incubation times, and ddNTP(s) for 3′ labeling. Significant differences were measured as described in **(B**).

We also investigated how the buffer used for working-stock enzyme storage at -20 °C impacted enzyme activity after prolonged periods of storage. We created working stocks by 5-fold dilution of a long-term -80 °C stock into a variety of different solution conditions and tested the activity of these working stocks. While many conditions did not make a difference in 3′-labeling or cDNA synthesis activity when assayed immediately after dilution, differences emerged after months of storage time that could be detected by assays of 3′-labeling activity directly or by production of OTTR cDNA libraries. We found that the following improved stability of BoMoC 3′-labeling activity during 6 months of storage at -20 °C (data not shown): mildly acidic (pH 6 – 6.5) buffers such as Bis-Tris (25 mM) or arginine-HCl (0.5 M); either increasing the concentration of dithiothreitol (DTT) (5 – 15 mM) or replacing DTT with a low concentration of Tris(2-carboxyethyl)phosphine (TCEP) (0.2 mM); and replacing KCl (800 mM) for a mix of KCl (200 mM) and (NH_4_)_2_SO_4_ (400 mM). A strong correlation emerged between conditions that would allow oxidation of the reducing agent and poor 3′-labeling activity. Maintaining the presence of reduced DTT by lowered pH or higher initial DTT concentration, or using a low concentration of pH-stable TCEP instead, were all effective for reducing loss of 3′-labeling activity (**Fig. 3B**, **Table 1**).

**Table 1:**
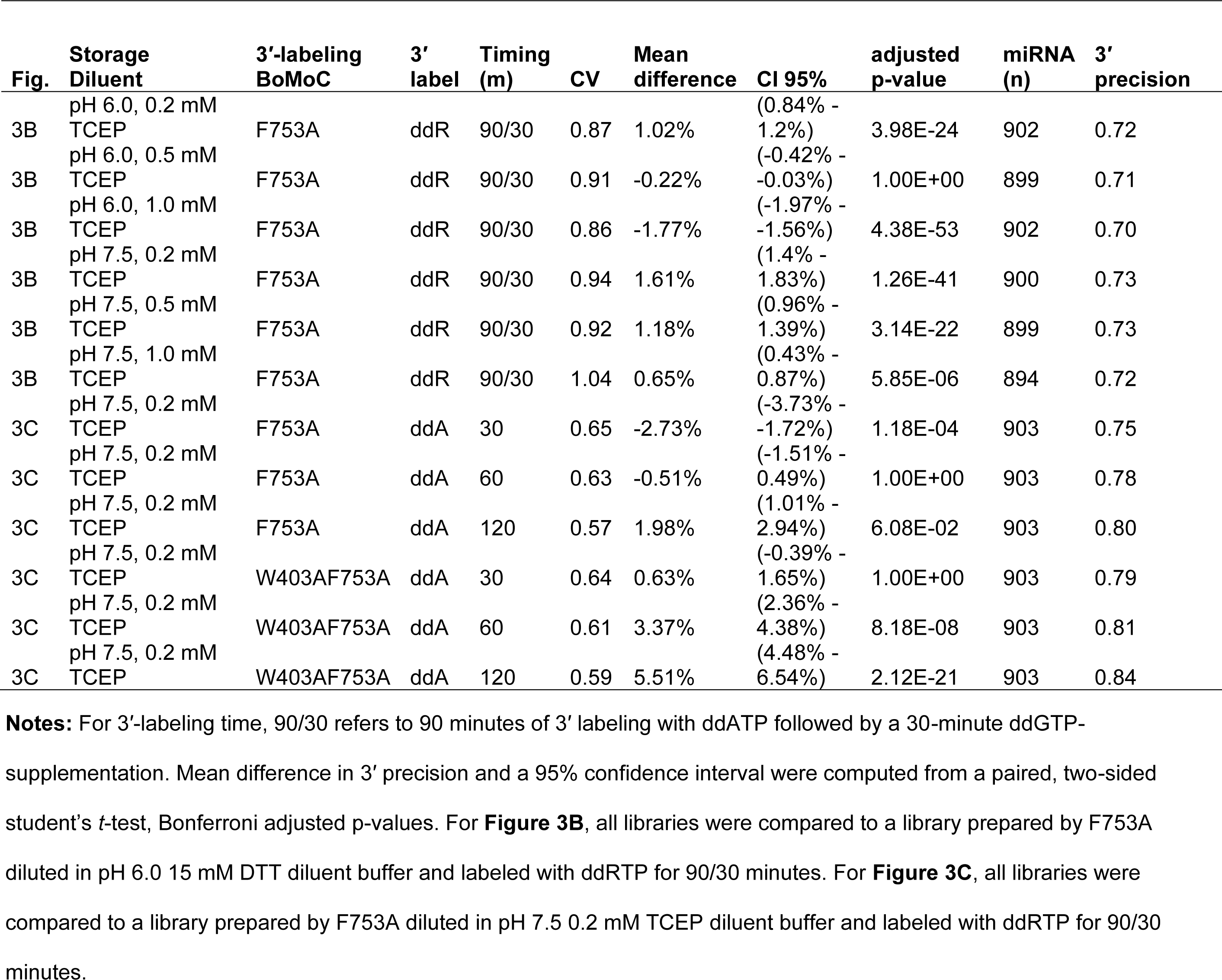
Impact of TCEP in enzyme storage diluent on 3′-labeling activity.

Given the results above, and a concern that excessively high DTT concentration could inhibit the phosphatase activity of the rSAP enzyme added after 3′ labeling (**Fig. 1**), we settled on an updated -20 °C storage diluent with neutral pH and low TCEP: 25 mM Tris-HCl pH 7.5, 200 mM KCl, 400 mM (NH_4_)_2_SO_4_, 0.2 mM TCEP, 50% glycerol (**Fig. 1**, changes in red). We also detected that over time a second OTTR buffer was sensitive to prolonged storage. After multiple free-thaw cycles within a 1 – 3 month time frame, the manganese in 3′ labeling Buffer 1B (**Fig. 1**) could form an oxidized yellow precipitate, which correlated with reduced 3′-labeling activity and thus reduced 3′ precision. This oxidation was defeated by exploiting prior knowledge (Hem 1963) to guide testing of replacement Buffer 1B, resulting in adoption of Buffer 1B2 with 10 mM sodium acetate at pH 5.5, 28 mM MnCl_2_, 28 mM (NH_4_)_2_SO_4_, and 3.5 mM ddATP (**Fig. 1**, changes in red).

With the 3′-labeling step optimized by BoMoC mutagenesis and improved enzyme and buffer storage stabilities, we assessed whether 3′ precision still required the 30-minute ddGTP 3′ labeling chase after 90 minutes of 3′ labeling with ddATP (**Fig. 1**). This ddGTP supplementation suppressed the over-capture of a small fraction of RNAs that did not 3′ label with ddATP under earlier OTTR conditions (Upton et al. 2021, Ferguson et al. 2023). Capture of 3′ddG-labeled input RNA obliged the +1 T primer duplex to be spiked with a lower concentration of +1 C primer duplex, where the 3′T nucleotide was changed to 3′C, a mutation that would compromise Read2 sequencing (**Table 2**, +T.v2.Cy5 and +C.v2.Cy5). If commercially available indexing primers that included the full-length Read2 sequence were used for library PCR, the cDNAs captured by the +1 C primer duplex would be under-amplified (**Supplemental Fig. 2)**. While these sequencing and amplification concerns could be readily overcome by adding an extra primer-duplex base pair (**Table 2**, updated universal primers for +1 Y duplex), we hoped to eliminate the ddGTP supplementation step entirely.

**Table 2:**
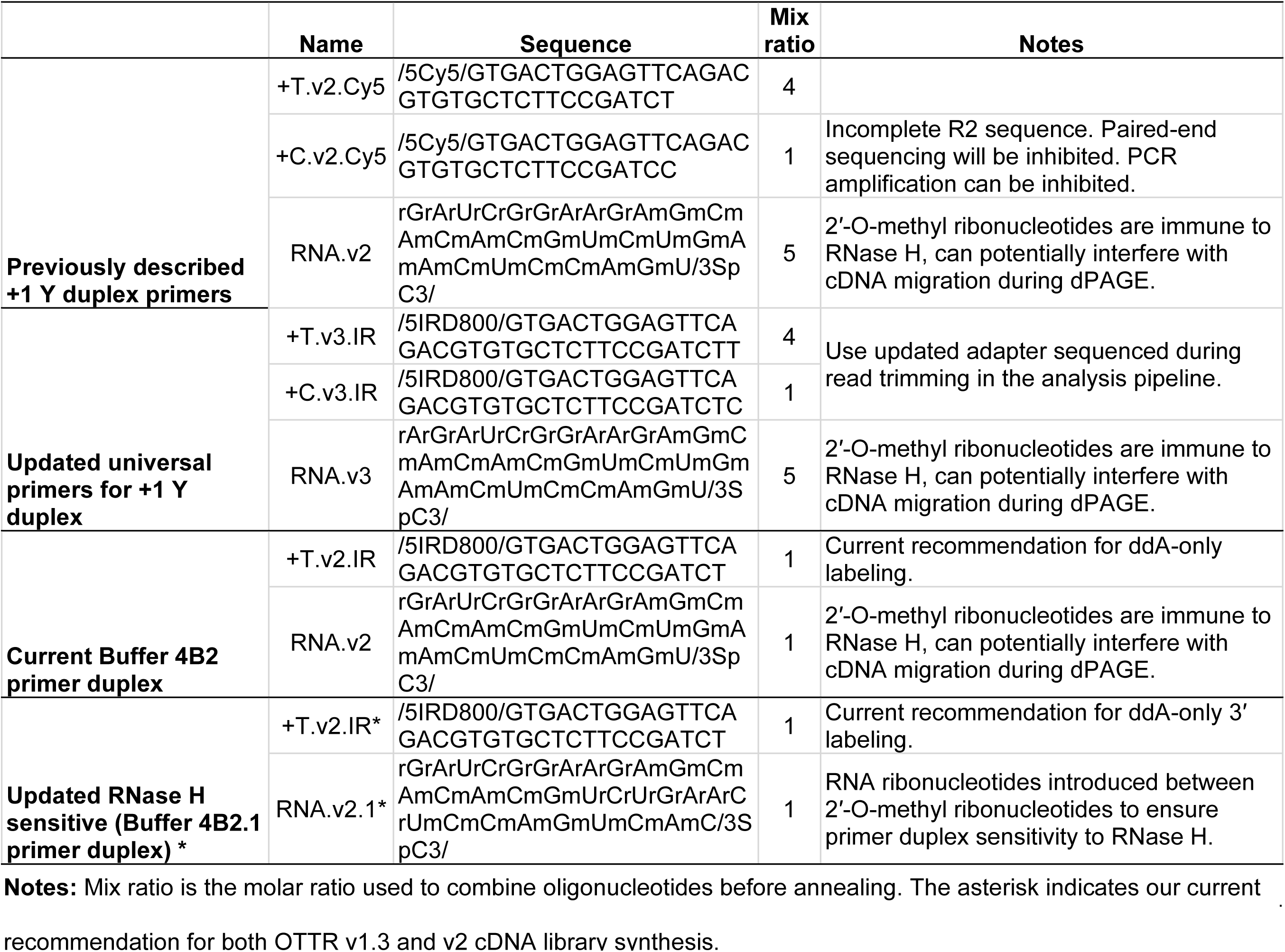
Primer duplex oligonucleotides used to capture ddA +/-3′ddG 3′-labeled input RNA.

To reassess 3′-labeling efficiency, we conducted time-course experiments under various conditions. While 30 minutes of 3′ labeling with ddATP by BoMoC F753A gave significantly worse 3′ precision than the standard OTTR protocol of 90 minutes of 3′ labeling with ddATP by BoMoC F753A followed by a 30-minute ddGTP supplementation, both 60 and 120 minutes of 3′ labeling with ddATP by BoMoC F753A were not significantly different for 3′ precision (**Fig. 3C**; **Table 1**). For BoMoC W403AF753A, we noted a significant improvement for ddA-only 3′ labeling at both 60 and 120 minutes of labeling compared to the standard OTTR protocol (**Fig. 3C**, **Table 1**). Moreover, we found the use of BoMoC F753A or W403AF753A for 120 minutes of 3′ labeling with only ddATP improved CV (0.568 and 0.597, respectively) compared to the inclusion of a ddGTP-supplementation step (0.766) (**Fig. 3C**; **Table 1**). In fact, all libraries synthesized with BoMoC F753A or W403AF753A using just ddATP showed markedly improved CV (0.568 – 0.647 or 0.597 – 0.646, respectively, **Table 1**).

### Precision of sequence capture at input RNA 5′ ends

Following cDNA synthesis across an input-template molecule, the cDNA is extended by a +1 G due to both a deoxyguanosine triphosphate (dGTP) skew in the dNTP concentrations used for the RT reaction and an inherent BoMoC preference for NTA using purine nucleotides (Upton et al. 2021). The NTA +1 G directs the second template jump to the 3′rC of the 3′ adapter template rather than another 3′ddR-labeled input template (**Fig. 1**). However, due to the high concentrations of adapter oligonucleotides in OTTR reactions, short cDNAs can be synthesized that are adapter dimers, generated by the side-reaction of a single template-jump from the +1 Y primer duplex to the 3′rC adapter template. Sanger sequencing of these short cDNAs cloned into a plasmid vector (**Table 3**) revealed that cDNAs with 3′ truncations of adapter template sequence were common, suggesting that the six nucleotides of RNA at the 3′ end of the original 3′ adapter template (Upton et al. 2021) were chemically labile and/or favored internal initiation. Also, in high-throughput sequencing of miRXplore cDNA libraries generated by both OTTR template-jumps, single-nucleotide miRNA 5′ truncations could be detected, suggestive of 5′ imprecision in sequence capture (**Fig. 4A**, **ii**).

**Figure 4:**
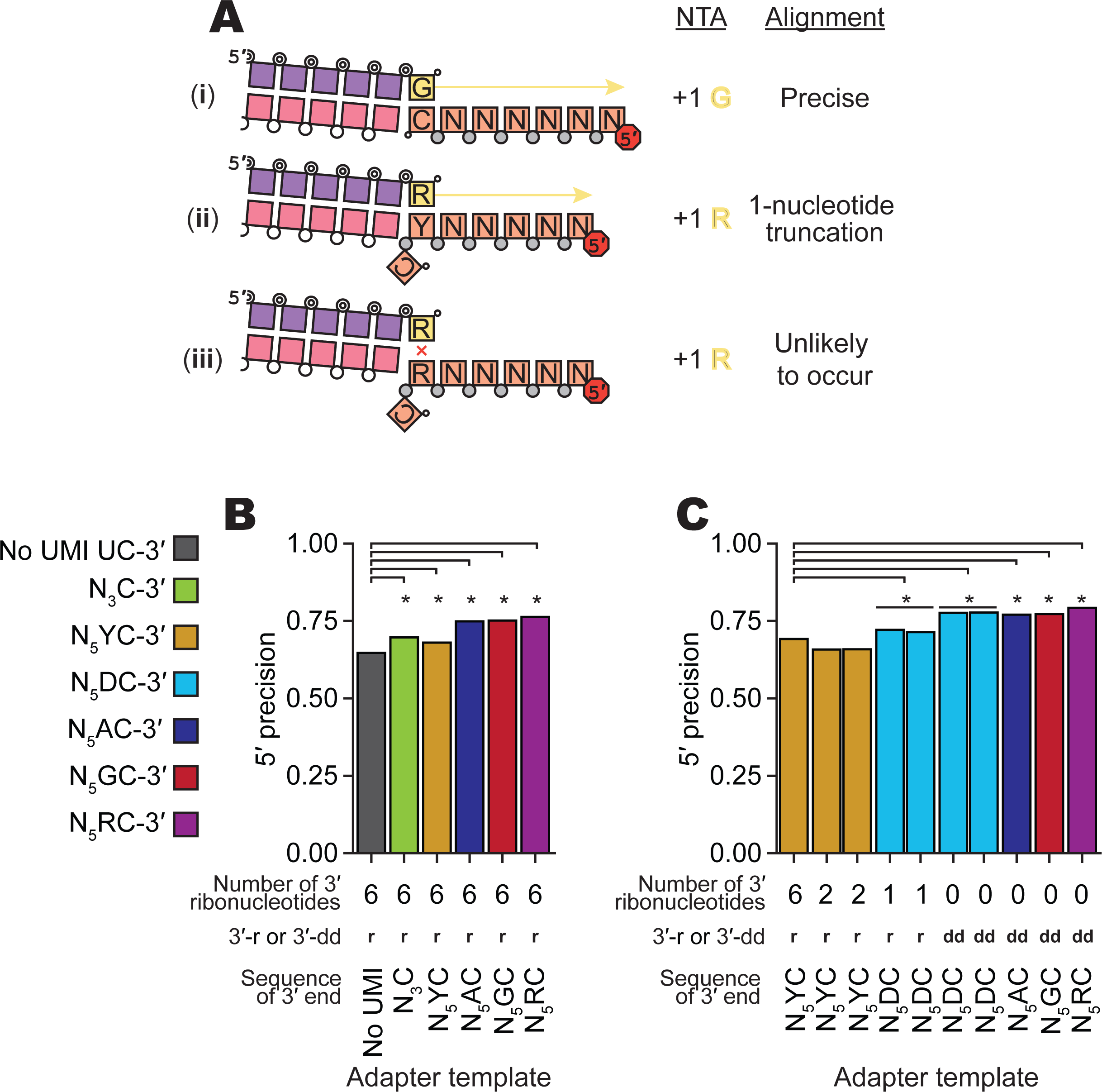
Evaluation of imprecision of end-to-end sequence capture at RNA 5′ ends. **A**, Mechanisms, numbered by Roman numeral, of precise and imprecise template-jumping to the 3′rC adapter template. Column one illustrates two possible and one unlikely specificity of sequence-junction formation following the color legend of Figures 1 and **2A**. Phosphodiester bonds are represented by circles: gray for chemically synthesized bonds, concentric double circle for BoMoC-polymerized, unfilled for input template RNA. Strand 3′ end symbols are a small black circle for dideoxynucleotide, a small open circle for 3′-OH, and a triangle for 3′-OH replacement with an unextendible carbon linker. The red octagon indicates the presence of a 5′ fluorescent dye. Column two defines the NTA that would prime 3′rC adapter-template capture. Column three defines whether the mapped template 5′-end was precise. **B,C**, Comparison of mean fraction of alignments with 5′ precision for libraries prepared with different 3′rC adapter template designs (**Table 5**). Significant differences in 5′ precision (denoted by asterisk) were determined by a paired, two-sided student’s *t*-test for each miRNA, with p-values adjusted by Bonferroni correction. See complete summary statistic in **Table 4**.

**Table 3:**
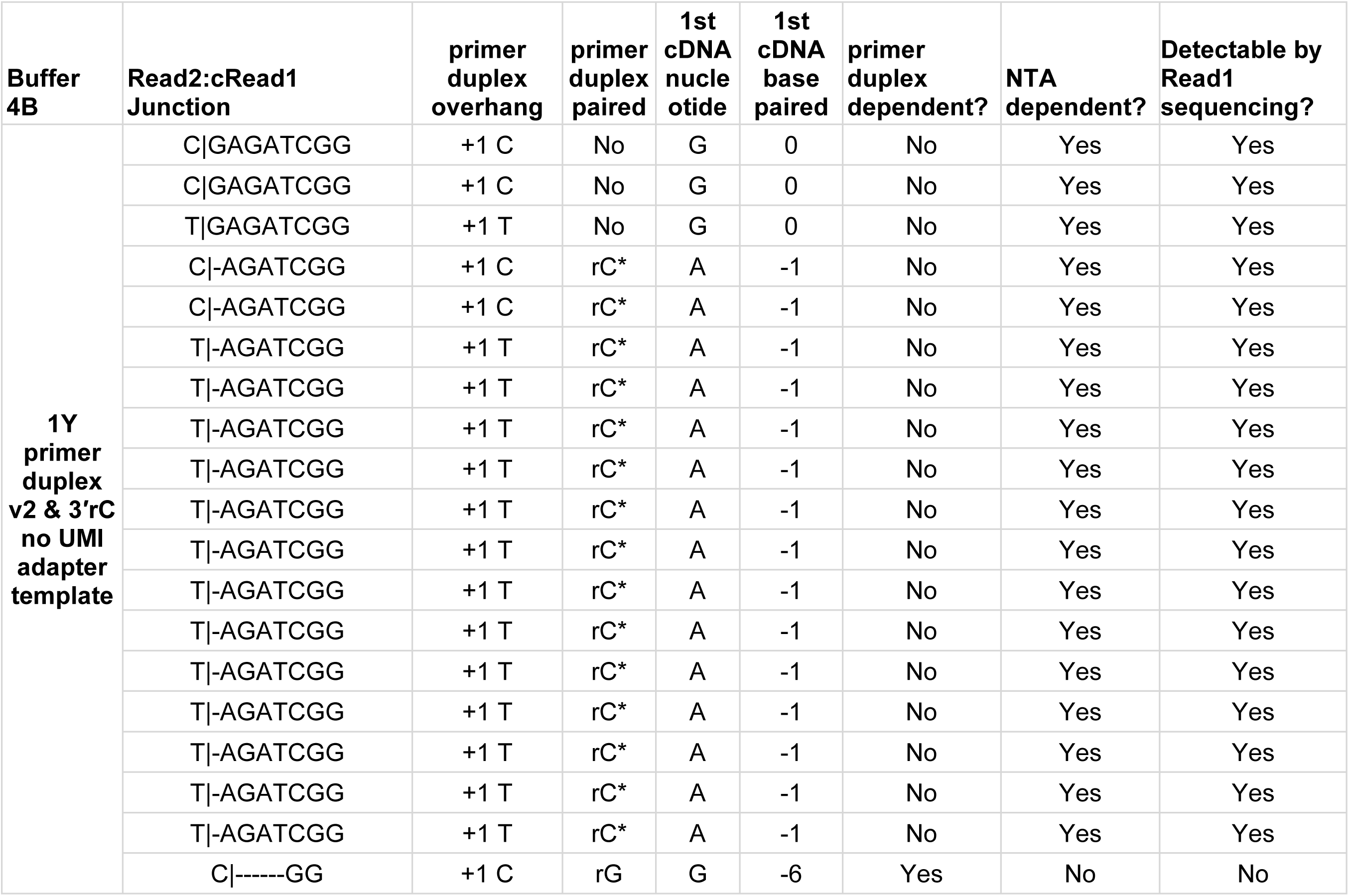
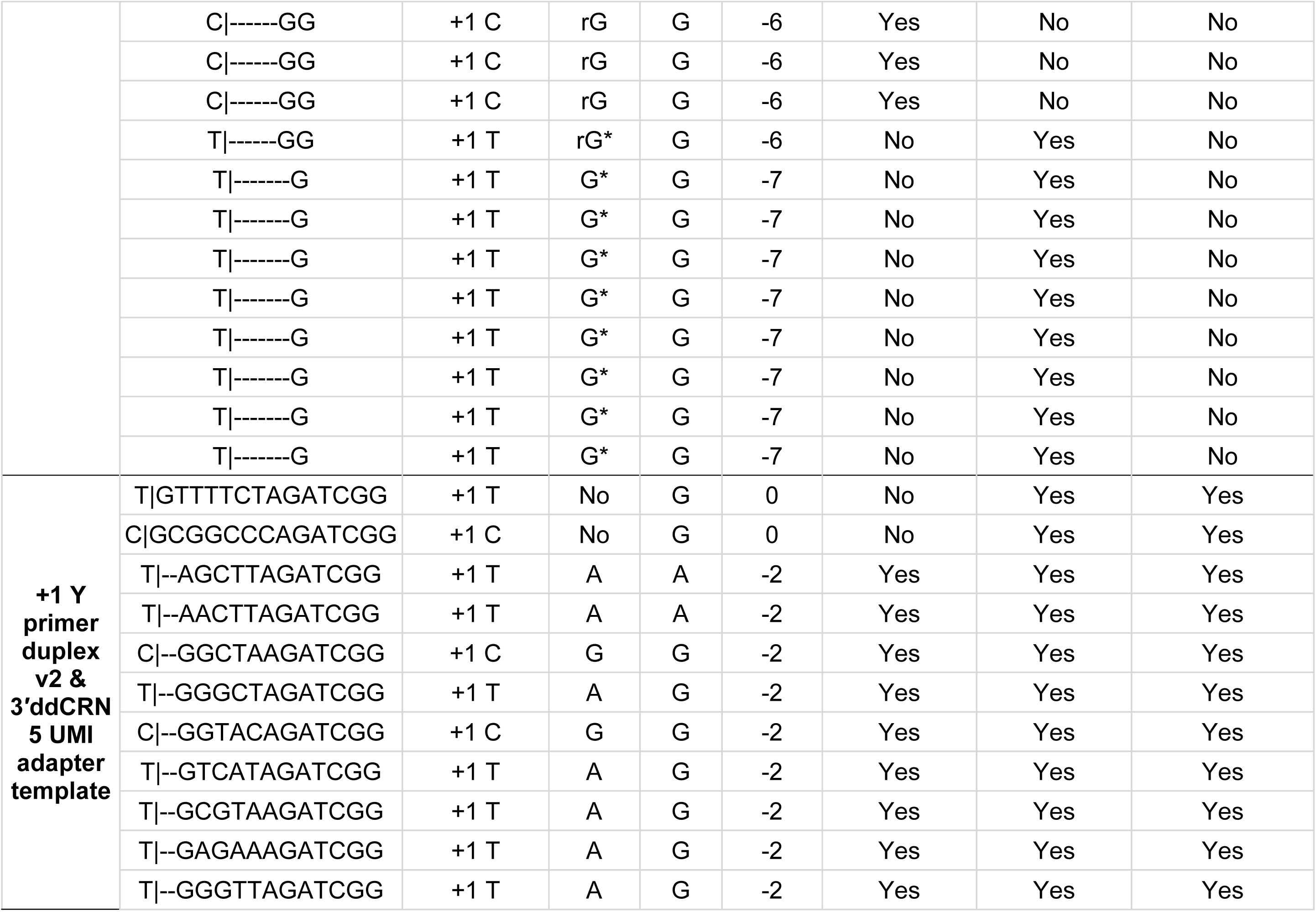
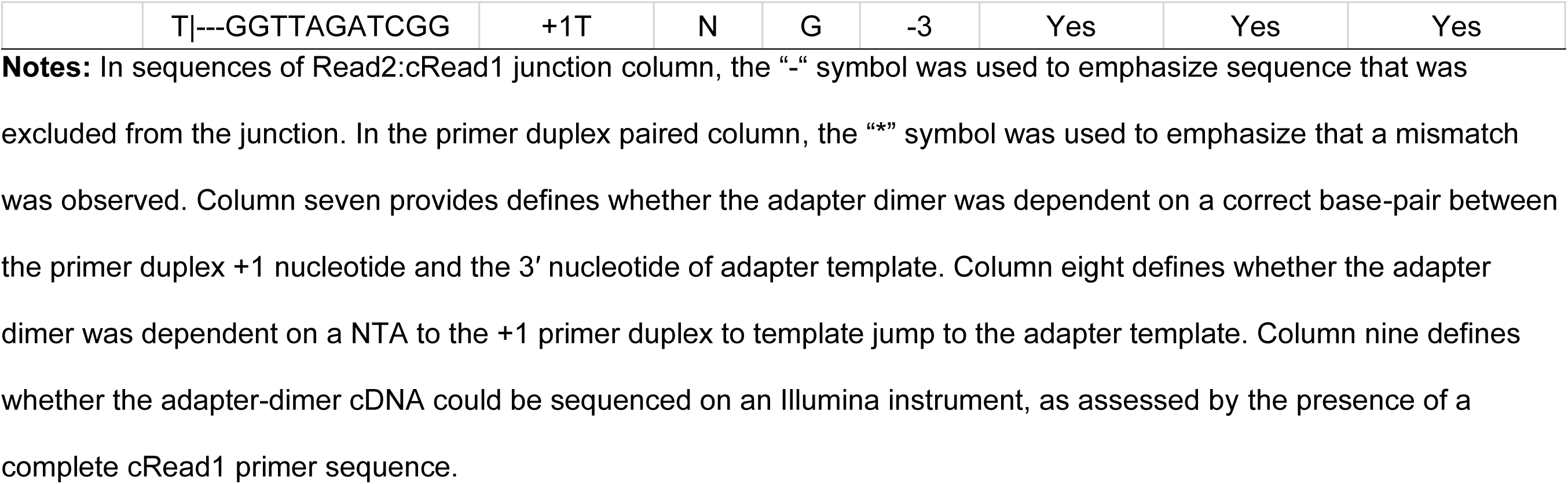
Summary of Sanger sequencing of plasmid-cloned adapter-dimer cDNAs.

**Table 4:** Impact of adapter template UMI sequence on complete 5′-end capture.

| Fig. | 3'r/ddC<br>adapter<br>template<br>sequence | 3'r/ddC<br>adapter<br>template<br># RNA | 3'r/ddC<br>adapter<br>template<br>end | CV | Mean<br>differen<br>ce | CI 95% | adjusted<br>p-value | miRNA<br>(n) | 5'<br>precision |
| --- | --- | --- | --- | --- | --- | --- | --- | --- | --- |
| 4B | NNNC-3' | 6 | 3'-OH | 1.07 | 4.92% | (4.64% - 5.21%) | 8E-160 | 902 | 0.7 |
| 4B | NNNNNYC-3' | 6 | 3'-OH | 0.65 | 3.3% | (2.99% - 3.62%) | 2.8E-74 | 903 | 0.68 |
| 4B | NNNNNAC-3' | 6 | 3'-OH | 0.72 | 10.1% | (9.56% - 10.65%) | 8E-176 | 889 | 0.75 |
| 4B | NNNNNGC-3' | 6 | 3'-OH | 0.73 | 10.45% | (9.98% - 10.91%) | 2E-221 | 883 | 0.75 |
| 4B | NNNNNRC-3' | 6 | 3'-OH | 0.81 | 11.54% | (11.04% - 12.03%) | 3E-232 | 881 | 0.76 |
| 4C | NNNNNYC-3' | 2 | 3'-OH | 0.66 | -3.4% | (-3.56% - -3.23%) | 4E-202 | 904 | 0.66 |
| 4C | NNNNNYC-3' | 2 | 3'-OH | 0.65 | -3.35% | (-3.51% - -3.18%) | 7E-196 | 904 | 0.66 |
| 4C | NNNNNDC-3' | 1 | 3'-H | 0.68 | 2.95% | (2.73% - 3.17%) | 1E-112 | 904 | 0.72 |
| 4C | NNNNNDC-3' | 1 | 3'-H | 0.69 | 2.24% | (2.02% - 2.46%) | 1.1E-72 | 904 | 0.71 |
| 4C | NNNNNDC-3' | 0 | 3'-H | 0.68 | 8.46% | (8.21% - 8.71%) | 0 | 904 | 0.78 |
| 4C | NNNNNDC-3' | 0 | 3'-H | 0.66 | 8.53% | (8.3% - 8.77%) | 0 | 904 | 0.78 |
| 4C | NNNNNAC-3' | 0 | 3'-H | 0.56 | 7.87% | (7.37% - 8.36%) | 2E-141 | 904 | 0.77 |
| 4C | NNNNNGC-3' | 0 | 3'-H | 0.61 | 8.06% | (7.62% - 8.49%) | 1E-175 | 903 | 0.77 |
| 4C | NNNNNRC-3' | 0 | 3'-H | 0.62 | 10.05% | (9.64% - 10.46%) | 2E-249 | 904 | 0.79 |

**Notes:** Figure panel and descriptions of the 3' adapter template (3' sequence, number of RNA nucleotides, and whether the 3' end was unblocked, *i.e.* 3'-OH, or blocked, *i.e.* 3'-hydrogen (3'-H)) are described in the first four columns. See **Table** **5** for complete adapter template sequences. The mean difference in 5' precision and a 95% confidence interval were computed from a paired, two-sided student's *t*-test, with Bonferroni adjusted p-values. For **Figure 4B**, all libraries were compared to a library prepared with no UMI in the 3'rC adapter template, and for **Figure 4C**, all libraries were compared to
a library prepared with 3'rC adapter template with 6-RNA ribonucleotides and a 3'-OH with the following 3' sequence:
NNNNNYC-3'.

**Table 5:**
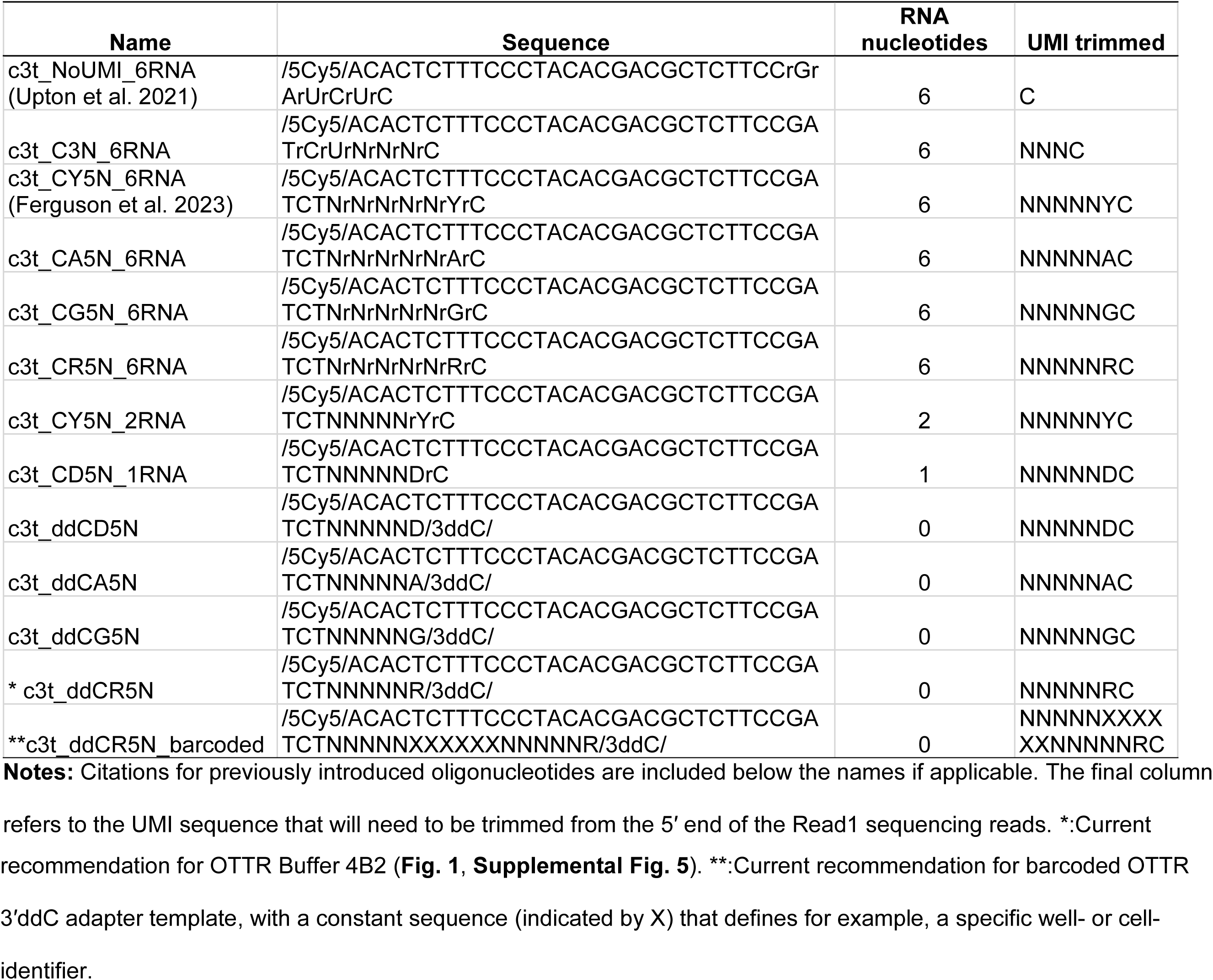
3′ adapter template oligonucleotide sequences.

To reduce 5′ imprecision, we evaluated the performance of different 3′ adapter-template sequences with specific and degenerate ribonucleotides between the 3′rC and Read1 sequence (see **Fig. 1** for schematic). In previous work (Ferguson et al. 2023), we included a UMI in the adapter template, with a 3′ penultimate rY followed by five degenerate nucleotides. Here, with improved understanding of capture imprecision at the input molecule 5′ end (the adapter template 3′ end), we hypothesized that an adapter template 3′ penultimate rR would discourage internal initiation whereas a 3′ penultimate rY would encourage it (**Fig. 4A, ii** versus **iii**). We first assessed 5′ precision using 3′ adapter-template sequences in the same format as the original 3′ adapter template, comprised mostly of DNA with RNA in the 3′-most six nucleotides, and found significant improvement in 5′ precision for all tested UMI-containing 3′ adapter templates (**Fig. 4B**, **Tables 4-5**). As predicted, 5′ precision was greatest using 3′ adapter templates with a purine nucleotide at the 3′-penultimate position (**Fig. 4B**; **Tables 4-5**).

We next tested how 5′ precision was affected by the 3′-terminal content of ribonucleotides versus deoxyribonucleotides in the adapter template. We replaced some or all of the 3′-terminal RNA with DNA, including a DNA-only 3′ adapter template with 3′ dideoxycytidine (ddC) (hereafter 3′ddC adapter template). Independent of adapter-template sequence, the replacement of RNA and inclusion of a 3′-terminal ddC improved 5′ precision (**Fig. 4C**; **Tables 4-5**). Strategically, the 3′ddC also prevented an adapter template from priming unwanted side-reaction synthesis. Of note, we did not detect an efficiency bias in comparison of jumping to DNA-only 3′ adapter template versus adapter templates with six 3′ ribonucleotides when assessing OTTR cDNA libraries produced using mixed populations of adapter templates, for example when adapter templates with six 3′ ribonucleotides and the sequence 3′rCrA were combined in a 1:2 ratio with the DNA-only 3′ddCG adapter templates (data not shown). Based on these findings, we adopted the use of a 3′ adapter template composed entirely of DNA with the terminal sequence 3′ddCR. This updated 3′ adapter template was included in the replacement of Buffer 4B with Buffer 4B2 (**Fig. 1**, changes in red).

### Lower limits for amount of input RNA

Optimization trials described previously (Upton et al. 2021) and above used 0.5 – 10 ng of miRXplore miRNA per OTTR reaction. To investigate OTTR performance across a wider range of input RNA amounts, we tested dilutions of the miRNA pool to span the range of 4 – 500 pg per 20 µL reaction, corresponding to 0.59 – 74 fmol input RNA. In an initial comparison, we used the previous best-practice conditions of BoMoC F753A 3′ labeling with ddATP for 90 minutes and a 30-minute additional 3′ labeling chase supplementation with ddGTP (Ferguson et al. 2023). As previously, the OTTR reaction used primer duplexes with +1 Y at a 4:1 ratio of +1 T to +1 C (**Table 2**, +T.v2 and +C.v2 annealed to RNA.v2), with the primer DNA strands bearing either the previous 5′ Cyanine5 (Cy5) or a 5′ near-infrared (IRD800) dye for fluorescence detection, and the previously standard c3t_CY5N_6RNA 3′rC adapter template (**Table 5**).

Detection sensitivity for cDNA resolved by denaturing (d) urea polyacrylamide gel electrophoresis (PAGE) was higher using IRD800 than Cy5 (**Fig. 5A**), and it was also possible to distinguish 5′-IRD800-labeled cDNAs from 5′-Cy5-labeled adapter template, which sometimes reannealed with cDNA during dPAGE (**Supplemental Fig. 3A**). To capture all cDNA products, cDNA size-selection encompassed an input nucleic acid size range of up to 200 nucleotides (**Fig. 5A**, black bracket). The sequenced libraries were assessed by the rank-order of their miRNA counts, which was consistent across the titration independent of what primer fluorophore was used (Spearman’s ρ, **Fig. 5B**). Library-wide CV of the miRNA count varied slightly across the titrations, with higher inputs yielding better (*i.e.,* lower) CVs (**Fig. 5B**). At all titration points, library-wide CV was below the original protocol CV of 0.86 (Upton et al., 2021).

**Figure 5:**
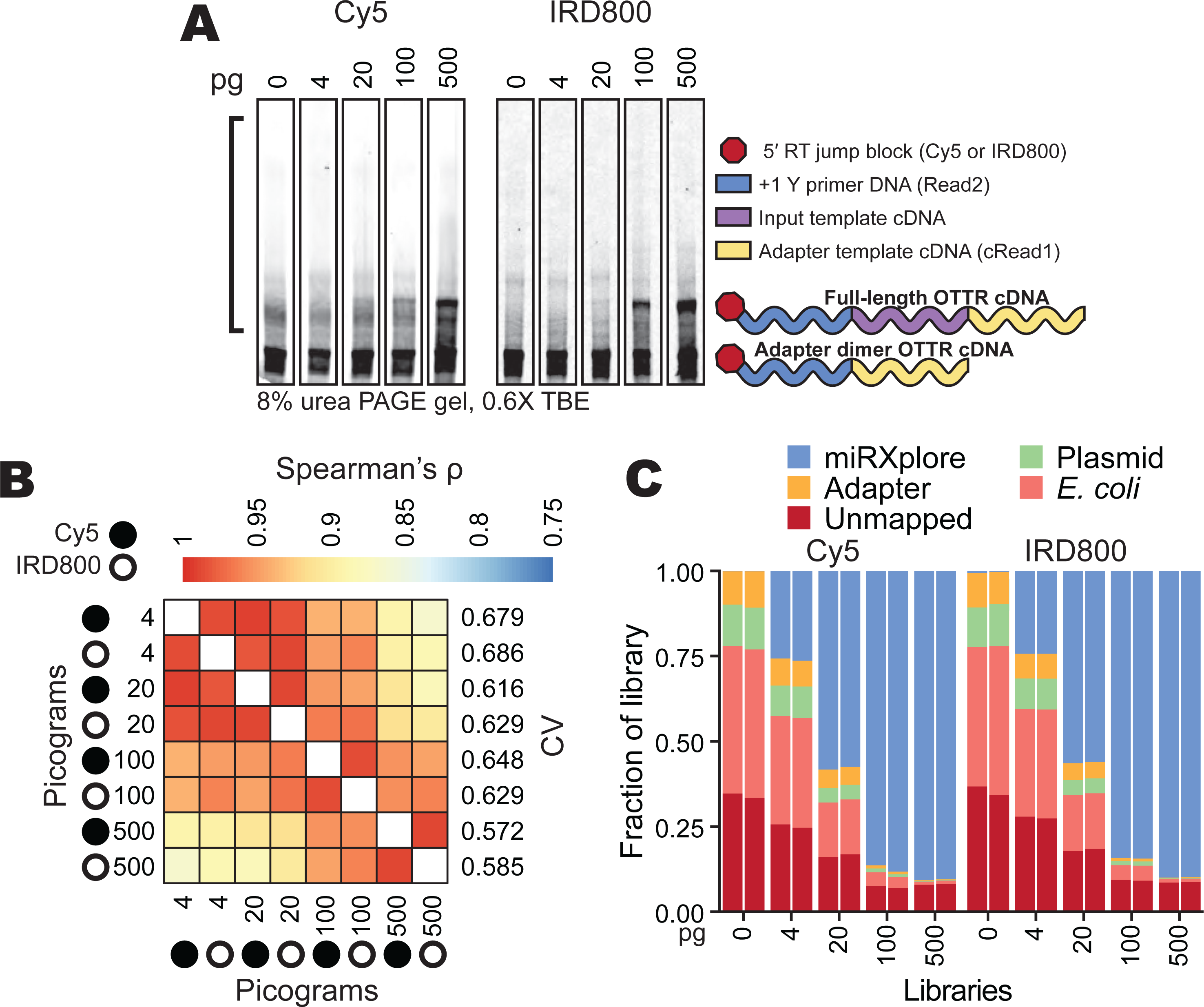
OTTR performance across different amounts of input RNA. **A**, Representative replicates of OTTR cDNA resolved by 8% dPAGE and detected using the 5′ Cy5 or IRD800 primer fluorophore, with miRXplore miRNA input in pg and 5′ primer fluorophore denoted above. cDNA product size-selection range is denoted by the left open bracket (black). On the right, adapter-dimer and miRXplore OTTR cDNA products are indicated with schematics using the color key from Figure 1. **B**, Correlation matrix of miRNA read counts from cDNA libraries produced using a titration from 4 to 500 pg total miRXplore RNA. Replicate Cy5 and IRD800 libraries for each miRNA input amount were averaged by their counts per million reads (CPM) and compared by Spearman’s correlation coefficient (ρ). miRXplore miRNA input in pg, 5′ primer fluorophore, and library-wide CV are denoted. **C**, Composition bar plots of the fraction of the total library, excluding reads 17 nucleotides or shorter, that mapped to miRXplore miRNA, OTTR adapter sequences, BoMoC expression plasmid, or *E. coli* genome, or were unmapped. Individual replicate libraries are presented in pairs.

While library-wide CV was consistently low across the input titration (0.572 – 0.686), the percentage of library with useful sequencing reads was not. Non-miRNA reads increased as input decreased (**Fig. 5C**). Most non-miRNA reads mapped either to *E. coli* nucleic acids or BoMoC expression plasmid (**Fig. 5C**, **Supplemental Fig. 3B**), both of which would be brought to the reaction by purified BoMoC. Because the same reaction conditions were used across the input titration, including a fixed amount of each enzyme, the relative representation of bacterial contaminants increased with the decrease in input template. We conclude that detection sensitivity can be limited by the competing capture of nucleic acid impurities in the enzyme preparations, despite rigorous discrimination against this in development of the original purification (Upton et al. 2021).

To increase the productive sequencing of low-input samples, we tested changes to the 3-step BoMoC purification regimen (**Supplemental Fig. 4**). Original purifications used Rosetta2 (DE3) pLys cells, which over-express seven rare-codon *E. coli* tRNA genes from a plasmid. BoMoC protein production was induced overnight at 16 °C, followed by cell lysis in the presence of benzonase to degrade DNA and RNA, lysate clarification, and chromatography using nickel, heparin, and size-exclusion chromatography (SEC) in series (**Supplemental Fig. 4**) (Upton et al. 2021). We screened an initial panel of enzyme purification changes by production of sequencing libraries using 20 pg of miRXplore as OTTR input and size-selection for cDNA inserts from ∼10 – 200 nucleotides as above. Comparison of two different enzyme purification strategies, starting from the same bacterial cell culture, revealed a change in the contaminating *E. coli* nucleic acids when polyethyleneimine (PEI) was used to clarify the cell lysate prior to chromatography (**Fig. 6A**, compare prep 2 that included a PEI precipitation step to prep 1 that did not). Sequencing reads that mapped to the BoMoC expression plasmid were greatly reduced, as were other *E. coli* nucleic acids, but full-length tRNAs remained (**Fig. 6B-C**). The level of tRNA contamination varied from purification to purification (data not shown), which we speculate resulted from slight differences in cell growth or lysis conditions for Rosetta2 (DE3) pLys cells.

**Figure 6:**
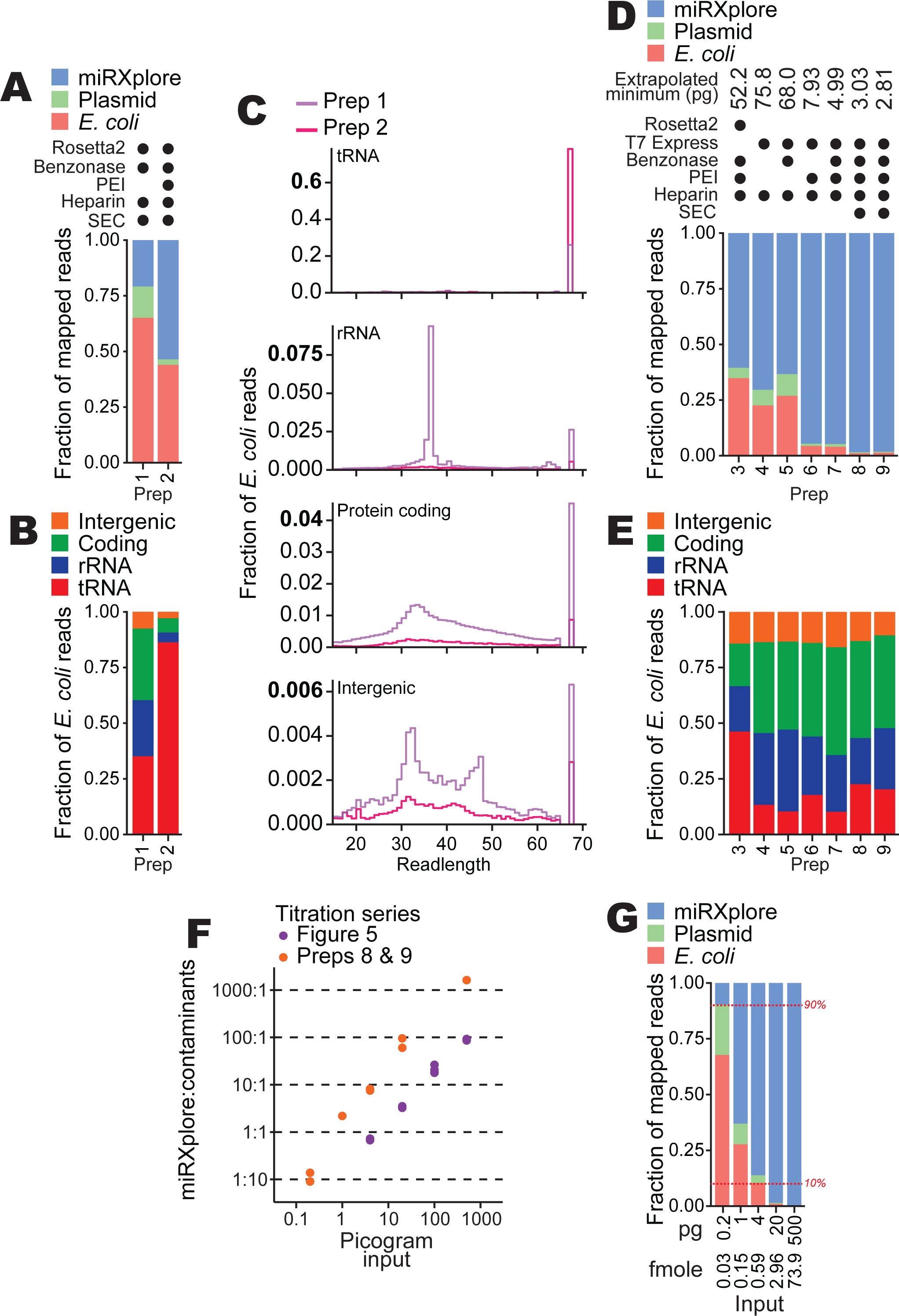
Improvements in BoMoC protein purification. **A**, Composition bar plots of mapped reads, excluding reads 17 nucleotides or shorter and adapter-mapping reads, for miRXplore cDNA libraries made using 20 pg input RNA and different protein purifications (preps). Variations to protein purification (**Supplemental Fig. 4**) are described using filled circles to indicate the presence of a variable *E. coli* strain or purification step in each prep. Prep 1 and 2 were BoMoC W403AF753A and were purified after splitting the same cell lysate in half. **B**, Subcategories of read mapping for the *E. coli* nucleic acid category in (**A**). **C**, Read length distribution plots of for the subcategories of read mappings in (**B**). Note that the y-axis for each plot has a different scale. **D,E**, Composition bar plots as described for (**A-B**) for cDNA libraries from 20 pg miRXplore input. In addition to 20 pg miRXplore RNA input, a 500 pg miRXplore RNA input cDNA library was produced; the pair were used to extrapolate minimum input that would recover 9:1 miRNA:*E. coli* and expression-plasmid reads, with that amount indicated at top. Preps 3 and 8 were BoMoC F753A; preps 4, 5, 6, and 7 were BoMoC W403AF753A; and prep 9 was BoMoC WT. **F**, Titration results for data from Figure 5 and additional results for cDNA libraries prepared using preps 8 and 9 for 3′ labeling and cDNA synthesis, respectively. The y-axis shows the read count ratio of miRXplore:contaminants (*i.e., E. coli* and expression plasmid sequences). Both axes are on a log_10_ scale. **G**, Composition bar plots as described for (**A**), here for libraries constructed with 0.2, 1, 4, 20, and 500 pg of miRXplore RNA and using preps 8 and 9 for 3′ labeling and cDNA synthesis, respectively. One representative replicate from the data in (**F**) was used for the bar plot. Red horizontal dashed lines (10% and 90%) are included as visual aids.

To quantify the extent of nucleic acid contamination across different enzyme purifications, we established an approach to estimate the minimum miRXplore input needed for ∼90% of mapped sequencing reads to map to miRXplore versus ∼10% to *E. coli* or BoMoC expression plasmid. OTTR libraries with inputs of 20 or 500 pg of miRNA were produced, sequenced, and mapped to reference sequences in parallel. Purifications compared the use of the Rosetta2 strain to an *E. coli* expression strain that does not over-express tRNAs, T7 Express lysY/Iq. Purifications also compared inclusion or omission of benzonase in the cell lysis buffer, lysate clarification with PEI, and the final SEC step. Comparison of read mapping using the 20 ng input samples revealed that the change of expression strain and the use of PEI combined were sufficient to nearly eliminate contaminating bacterial nucleic acids, with some additional benefit when SEC was included (**Fig. 6D**). Purifications of BoMoC F753A and BoMoC WT from T7 Express cells lysed in the presence of benzonase, and with cell lysate clarified by PEI precipitation (preps 8 and 9, respectively, in **Fig. 6D**), gave the lowest level of contaminating nucleic acid reads, which showed reduced predominance of *E. coli* tRNA relative to *E. coli* ribosomal RNA (rRNA) and coding- or intergenic-region of the *E. coli* genome (**Fig. 6E**).

We used OTTR cDNA libraries produced from a range of miRXplore input amounts to extrapolate the input requirement for 90% or more of reads to map to input templates instead of contaminant bacterial nucleic acids. From the input RNA titration series described above performed using BoMoC purified with the original protocol, the input amount required for reads to be 90% miRXplore versus bacterial nucleic acids was 52 pg (**Fig. 6F**, purple data points). In comparison, using the optimized expression and purification conditions (**Fig. 6D**, preps 8 and 9), the input amount required for reads to be 90% miRXplore versus bacterial nucleic acids was only 2.8 pg (**Fig. 6F**, orange data points). We confirmed this extrapolated lower limit on input amount by generating OTTR cDNA libraries using a titration of miRXplore input from 0.2 – 500 pg in reactions using optimally purified proteins (**Fig. 6G**). We conclude the updated BoMoC purification protocol (**Supplemental Fig. 4**, enzyme purification protocol changes in red) substantially reduces the co-purification of bacterial nucleic acids, which is enabling for particularly low-input OTTR cDNA libraires. Lowering the BoMoC enzyme amounts used in low-input OTTR reactions would further decrease the minimum input for only 10% or less of reads to be non-productive bacterial nucleic acid sequencing.

### Enabling gel-free OTTR library synthesis

A remaining challenge for OTTR, as with other small RNA capture methods, is to limit production or improve removal of adapter-dimer cDNAs, which are sequenced at the expense of informative cDNAs. For many applications, for example ribosome profiling (Ferguson et al. 2023), the presence of adapter-dimer cDNAs necessitates OTTR cDNA size-selection by dPAGE (see **Fig. 5A**). This is a limitation for high-throughput automation of OTTR library production and for laboratories that rely on commercially manufactured dPAGE gels, which become less denaturing over time and therefore may fail to prevent reannealing of adapter oligonucleotides to OTTR cDNA during electrophoresis, compromising accurate size-selection. Approaches to reduce sequencing reads from OTTR adapter-dimer products can take advantage of features that distinguish a single-jump adapter-dimer from a double-jump, bona fide OTTR cDNA. One obvious difference is the fact that adapter-dimer cDNA duplexes contain only sequences derived from the primer duplex and adapter template, whereas desired OTTR cDNA duplexes contain a 3′-labeled input RNA.

Conveniently, a biotinylated nucleotide can be used in the input 3′ labeling reaction (**Fig. 7A**). Subsequently, after the OTTR cDNA synthesis step, the biotinylated input molecule can be used to selectively enrich the desired cDNA duplexes from adapter-dimer cDNA duplexes. Streptavidin enrichment of OTTR cDNA duplexes containing biotinylated ddATP was as effective as gel-based cDNA size-selection for removing adapter-only cDNAs (**Fig. 7B**). With low-input samples, the introduction of additional streptavidin washing steps would be prudent to more completely remove the remaining non-specifically captured adapter-dimer cDNA duplexes (**Fig. 7B**). The OTTR protocol version with 3′-labeling using biotinylated ddATP and subsequent cDNA duplex library enrichment by streptavidin (OTTR v2, **Supplemental Fig. 5**) reduces the need to suppress adapter-dimer formation. Therefore, we examined whether changing the cDNA synthesis conditions without concern for adapter dimer formation could reduce bias or increase precision of sequence capture. To this end, we screened alternative compositions for a Buffer 4A equivalent to be used in OTTR v2 (**Fig. 7C, Supplemental Fig. 5**).

**Figure 7:**
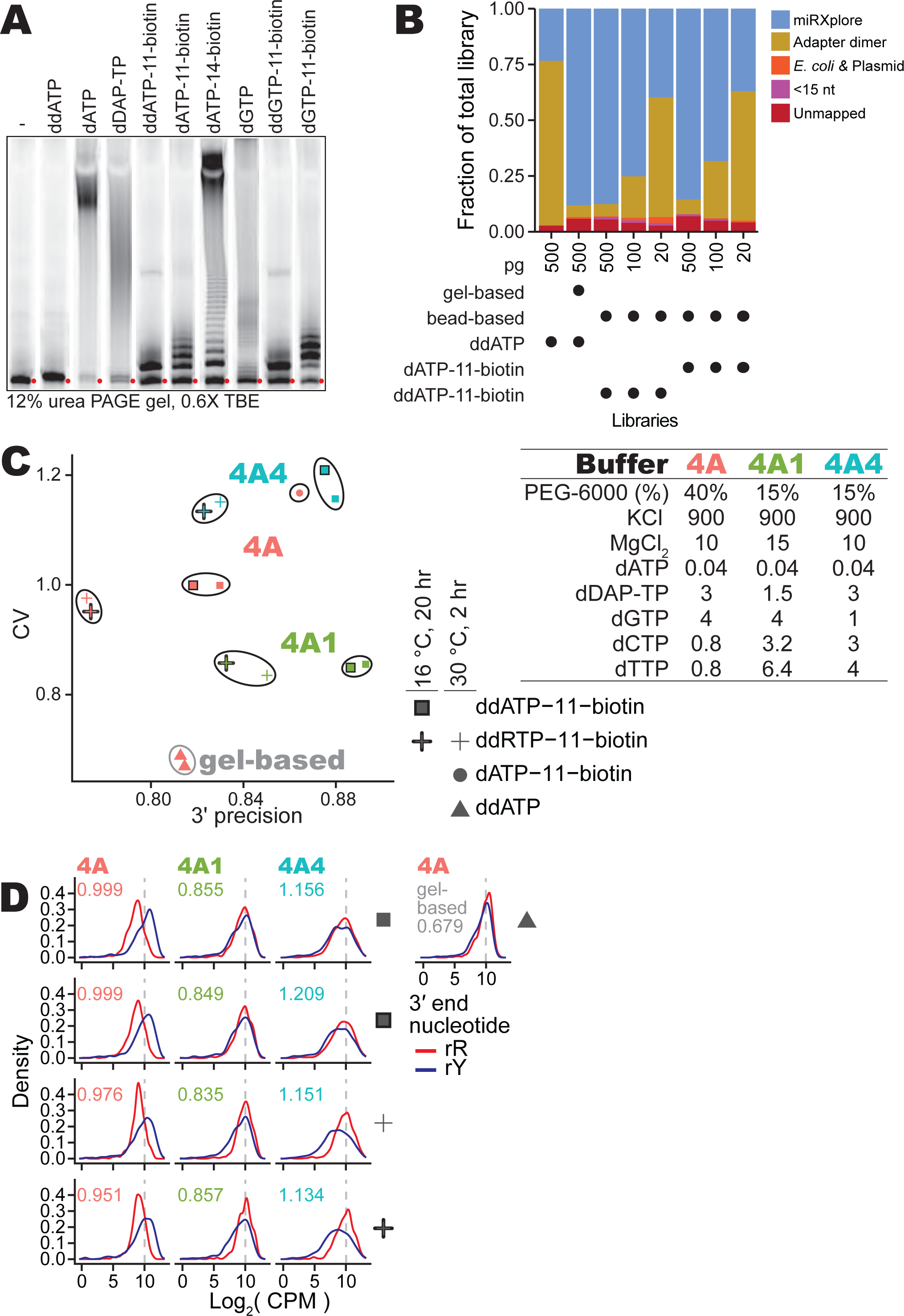
Gel-free OTTR by capture of biotinylated-input OTTR cDNA duplexes. **A**, The 3′ labeling activity of W403AF753A BoMoC with various d(d)RTPs resolved by 12% dPAGE and detected by the 5′ IRD800 fluorophore of the 3′NN single-stranded DNA substrate. The number in the biotinylated nucleotide names refers to the carbon linker. ddATP-11-biotin, ddGTP-11-biotin, dATP-11-biotin, and dGTP-11-biotin were biotinylated from the 7-position of the purine base while dATP-14-biotin was biotinylated from 6-position of the purine base. Red circles indicate the migration of primer without elongation. **B**, Fractional content of the total sequenced reads for different miRXplore miRNA libraries with different input amounts, 3′ labeling nucleotide, and post-cDNA synthesis clean-up (*i.e.,* either gel-based or streptavidin pull-down). **C**, Scatter plot of miRXplore miRNA library CV and 3′ precision of various biotinylated OTTR libraries. The Buffer 4A (either 4A, 4A1, or 4A4) used in each library is labeled by color of the data points. Buffer 4A recipes are specified at right. The shape of the data points indicates the nucleotide and reaction condition used during 3′ labeling. Libraries with the same 3′ labeling nucleotide and Buffer 4A are grouped together by a bounding line as a visual aid. **D**, Distributions of the log_2_ CPM of miRXplore miRNA based on miRNA 3′ nucleotide for the libraries in **C**, with red for purine and blue for pyrimidine as indicated by the key. Library-wide CV for each library is given in the top-left corner of the plots. Color of CV text indicates which Buffer 4A variant was used. A representative gel-based cDNA purification of OTTR cDNA library was included at right.

With the Buffer 4A used for OTTR v1.3 (**Fig. 1**), 3′ biotinylated templates were captured with a bias for miRNAs with 3′rY (**Fig. 7D**, 1^st^ column). The lowest bias and highest 3′ precision were attained by increasing dTTP and dCTP concentration while reducing diaminopurine deoxynucleotide triphosphate (dDAP-TP) concentration (**Fig. 7C**, Buffer 4A1; **Fig. 7D**, 2^nd^ column). Increase in 3′ precision was observed using elevated dTTP and dCTP alone, whereas decrease in library CV required the combination of elevated dTTP and dCTP and a reduction of dDAP-TP (**Fig. 7C**, compare results for Buffer 4A1 with results for Buffer 4A4; **Fig. 7D**, compare 2^nd^ to 3^rd^ column). We note that alternative 3′-labeling conditions, such as using both ddATP-11-biotin and ddGTP-11-biotin or using a lower reaction temperature combined with longer reaction time, did not suppress the preferential capture of 3′-rY miRNAs (**Fig. 7C-D**). We adopted Buffer 4A1 in the final optimized workflow for gel-free, automation-compatible OTTR v2 (**Supplemental Fig. 5**).

Using OTTR v2, we converted a pool of input RNAs to OTTR cDNAs, indexed the cDNAs by PCR, and loaded the libraries on an Illumina MiniSeq in a single day without any gel- or column-based clean-up or size-selection. The optimized OTTR v1.3 gel-based size-selection protocol (**Fig. 1**) gave modestly better uniformity of miRNA capture but less 3′ precision than the optimized gel-free OTTR v2 protocol (**Fig. 7C**, compare red triangles to green squares). The optimized OTTR protocols established from the work described here are provided as **Supplemental Method 1** and **Supplemental Method 2**.

### Future improvements and expanded applications

OTTR is highly amenable to bespoke tailoring. The work described above illustrates the adjustable balance between more quantitative representation of input RNAs (lower library CV) versus higher 3′ precision of sequence capture. The former is critical to reliably profile sequence diversity in a complex sample (*e.g.* for liquid biopsy studies), while the latter serves applications in which knowledge of the precise 3′ nucleotide matters (*e.g.* for ribosome profiling and miRNA or tRNA end-processing studies). The work above also introduces the consideration of trade-off between the advantages of cDNA size-selection by gel purification (*e.g.* for economy of sequencing only an input size range of interest) versus binding to streptavidin resin (*e.g.* to be automation-compatible).

Many additional OTTR protocol variations are possible. For example, to restrict input nucleic acid capture to particular 3′-end nucleotide(s) (*e.g.,* 3′rG RNAs produced by RNase T1 cleavage), a custom-designed primer duplex can be used (for 3′rG RNAs, primer duplex with +1 C made from annealed +C.v3 and RNA.v3, **Table 6**). As another example, for input-template capture without an initial 3′-labeling step, a mixture of +1 primer duplexes can be used (**Table 6**, v3 +1 set). Because +1 G primer duplex with 3′ddC adapter template would efficiently generate adapter dimer, splitting the OTTR cDNA library synthesis reaction into successive, single template-jump reactions would be prudent. The first template-jump would synthesize cDNA across the input template using a BoMoC variant and/or reaction conditions that limit NTA (Upton et al. 2021, Pimentel et al. 2022). The second template-jump would be initiated by providing the 3′ddC adapter template in reaction conditions that support +1 G NTA. As a third example, different sequences of primer duplex or adapter template could be used. We previously used the complete lengths of Illumina sequencing adapters (∼70 nucleotides) rather than only the R1 and R2 portions (∼35 nucleotides) to enable sequencing-ready OTTR cDNA library synthesis without PCR (Upton et al. 2021). It will be of interest to test adapter sequences used in applications other than Illumina sequencing and to test different UMI lengths and/or sample-identifying barcodes to support single-cell applications (*e.g.,* **Table 5**, c3t_ddCR5N_barcoded).

**Table 6:**
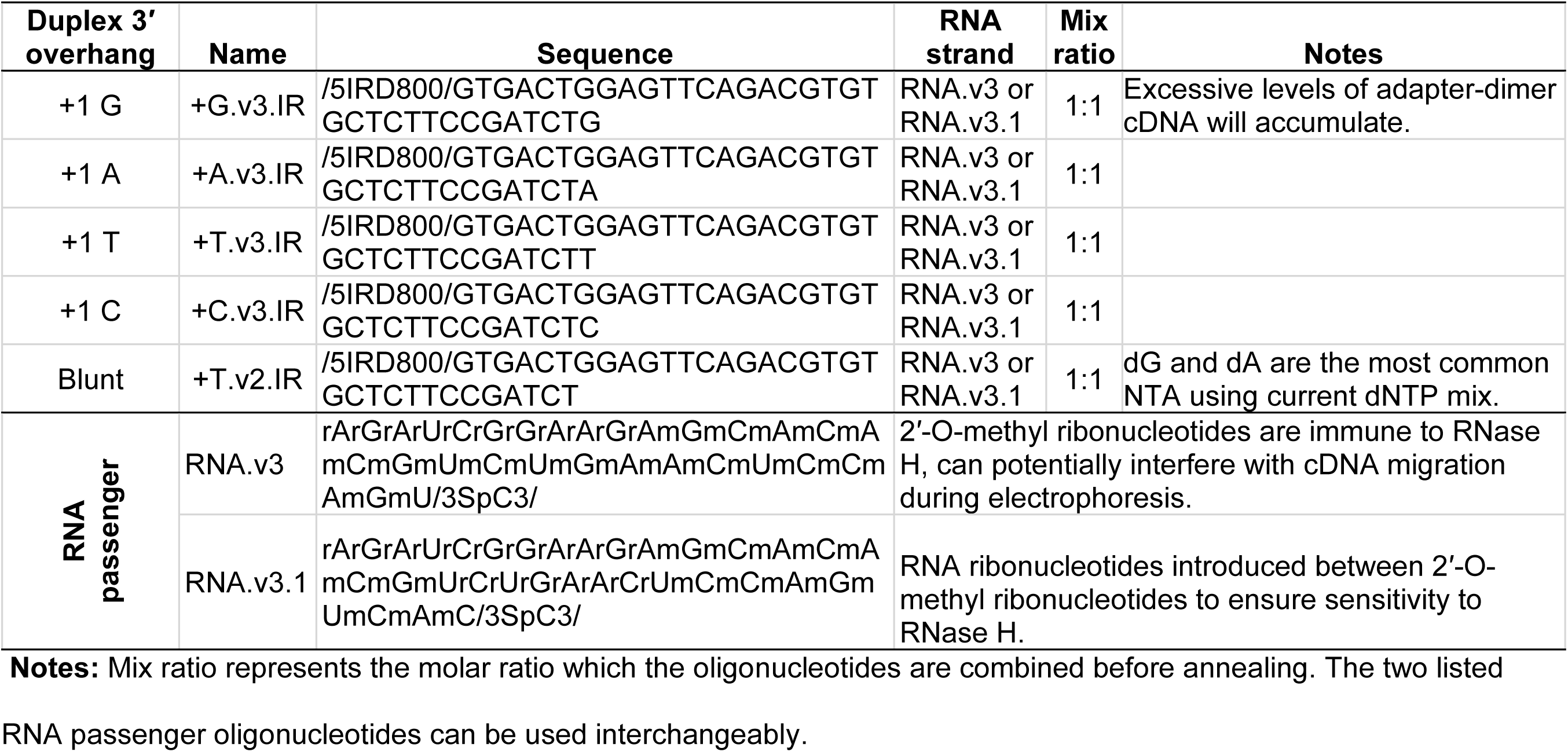
Alternative primer duplexes to be used for capture of RNAs with specific 3′ ends.

OTTR exploits non-canonical nucleotides in both the primer duplex RNA passenger-strand (**Table 2**, RNA.v2), which has several positions of RNA modification by duplex-stabilizing 2′-O-methyl groups (Upton et al. 2021), and in the 3′ddC DNA adapter template (**Table 5**, c3t_ddCR5N), which replaces the chimeric RNA-DNA adapter templates used in earlier versions of OTTR (Upton et al. 2021, Ferguson et al. 2023). One challenge in the use of these adapter oligonucleotides is that they can re-anneal with cDNA during dPAGE-based size-selection, which broadens the spread of adapter-dimer gel migration into the range of desired OTTR cDNAs (**Supplemental Fig. 3A**). One purpose of 2′O-Me substitutions in the primer-duplex passenger strand was to increase duplex stability, and another was to preclude use of the passenger strand as a template-jump acceptor. In the experiments of Figure 7, we used a primer-duplex passenger strand with ribonucleotide patches between blocks of 2′O-Me substitution to enable its fragmentation by RNase H (**Table 2**, RNA.v2.1, and **Table 6**, RNA.v3.1). In parallel, the 3′ddC DNA adapter template could be anchored at its 5′ end to a resin or plate and cDNA eluted by denaturation. The utility of these and other OTTR protocol variations would vary with the type of input RNA.

Of high future interest is the benchmarking of OTTR protocols optimized for use of input samples other than small cellular RNAs. OTTR should be suitable for production of sequencing libraries from forensic samples, ancient DNA, cell-free DNA, and formalin-fixed paraffin-embedded biopsy material, given the high tolerance of BoMoC for base-modified templates such as mature tRNAs (Gustafsson et al. 2022, d’Almeida et al. 2023, Manning et al. 2024, Davey-Young et al. 2024) and the low input requirement demonstrated above. OTTR remains to be optimized for these sample types and for lengths of nucleic acid longer than tRNA. Both RNA and DNA oligonucleotides can be efficiently 3′ labeled by BoMoC (Upton et al. 2021), but optimal accessibility of longer DNA or RNA 3’ ends, or 3’-labeling of RNA-hairpin templates like miRNA precursors, may require single-stranded binding proteins or other additives to suppress preferential input capture. Pre-denaturation of template structure may be advantageous before the 3′-labeling step and again before cDNA synthesis.

Finally, automation of sub-microliter scale reactions would enable OTTR use for single-cell library synthesis. A single human cell has an estimated ∼1.3 fg of miRNA in 10 pg of total RNA, roughly ∼115,000 molecules (Peltier and Latham 2008, Bissels et al. 2009). With OTTR reaction conditions used above, we extrapolate an input requirement of ∼230 cells to have less than 10% of mappable reads produced from *E. coli* nucleic acids in the standard 20 µl reaction. Because the enzyme amounts added to low-input OTTR reactions could be scaled down from the current protocol, single-cell miRNA sequencing seems feasible. Beyond miRNAs, the use of OTTR for single-cell ribosome profiling is equally worth considering. From 10 pg of total RNA in a human cell, assuming ∼9 pg rRNA, if that rRNA was entirely assembled into translating ribosomes, we calculate a theoretical maximum of 43.7 fg mRNA RPFs, ∼2,300,000 molecules, from a single cell. Even with current OTTR reaction conditions, we extrapolate an input requirement of only ∼7 cells to have less than 10% of mappable reads produced from *E. coli* nucleic acids. With optimization, single-cell surveys of miRNAs, tRNAs, and tRFs, or single-cell ribosome profiling using OTTR could be cost-effective and information-rich alternatives to current protocols for single-cell profiling (Vandereyken et al. 2023).

## Materials and methods

### Protein purification, -80 °C long-term storage, and -20 °C working stock

BoMoC enzymes were purified largely as described previously (Upton et al. 2021, Ferguson et al. 2023) using nickel affinity chromatography, heparin chromatography, and SEC, unless specified. Initial and improved protein purification conditions are compared in **Supplemental Figure 4**. BoMoC has a C-terminal 6x-histidine tag for purification, an N-terminal maltose binding protein fusion for solubility, and an inactivated endonuclease activity. Strains were Rosetta2 (DE3) pLysS (Novagen, 71403), which possesses a plasmid to over-express rare *E. coli* tRNAs, or T7 Express Lys/IQ (New England Biolabs, C3013I). Cells were transformed with an expression plasmid (*e.g.* Addgene plasmid 186461, 185710, or 185713) and cultured overnight in 100 mL of 2X yeast extract tryptone (YT) media supplemented with appropriate antibiotics. The next day, cultured cells were diluted to OD_600_ of 0.05 and incubated in a 37 °C orbital shaker until the OD_600_ was 0.5 – 0.6. The culture flasks were then transferred to a pre-chilled 16 °C orbital shaker and left to chill for 25 – 30 minutes. The OD_600_ was monitored until it reached 0.7 – 0.8, at which point isopropyl ß-D-1-thiogalactopyranoside was added to a final concentration of 0.5 mM to induce protein expression. After 12 – 16 hours, cells were harvested by centrifugation for 20 minutes at 3,600 x g at 4 °C. The media was aspirated from the cell culture pellet and replaced with 15 mL of lysis buffer (20 mM Tris-HCl pH 7.4, 1 M NaCl, 10% glycerol, 1 mM β-mercaptoethanol, 1 μg/ml pepstatin A, 1 μg/ml leupeptin, 0.5 mM phenylmethylsulfonyl fluoride) per L of culture. After resuspension in lysis buffer, the cell lysate was frozen in liquid nitrogen and stored at -80 °C.

Frozen cell lysate was thawed in a room-temperature water bath before supplementing with MgCl_2_ to 0.5 mM. If specified, a volume of benzonase (Millipore) was added corresponding to a 1:1000 dilution. The thawed lysate was transferred to ice and sonicated for 3.5 minutes with 10 seconds of sonication separated by 10 seconds of rest. Insoluble material was pelleted by centrifugation using a SS34 rotor at 15,000 RPM for 30 minutes at 4 °C. Supernatant was decanted and recentrifuged at 5,000 x g for 10 minutes at 4 °C to remove any additional insoluble material from the cell lysate supernatant. For preps where PEI precipitation was included, the cell lysate was decanted into a bottle with a stir bar and set to stir rapidly without foam. Prior to this, a 10% w/v PEI solution was made fresh, first by stirring 10 g PEI in 40 mL water while adding concentrated HCl dropwise until the pH was ∼7.2. Once cooled, the pH was adjusted to pH 7.0 – 7.4 and the volume was adjusted with additional water to to 50 mL total. The neutralized 10% w/v PEI solution was added dropwise to the cell lysate until a final concentration of 0.2% w/v PEI was reached. Nucleic acids were pelleted by centrifugation using a SS34 rotor at 15,000 RPM for 30 minutes at 4 °C, and the supernatant was decanted as clarified cell lysate.

All chromatography steps were at room temperature. For nickel affinity chromatography, clarified lysate was loaded to two 5 mL HisTrap FF Crude columns (Cytiva) connected in series. Unbound protein was removed by 5 – 10 column volumes of Nickel A buffer (20 mM Tris-HCl pH 7.4, 1 M KCl, 20 mM imidazole, 10% glycerol, 1 mM β-mercaptoethanol). Bound protein was eluted with five column volumes of Nickel B buffer (20 mM Tris-HCl pH 7.4, 1 M KCl, 400 mM imidazole, 10% glycerol, 1 mM β-mercaptoethanol). Fractions were measured for protein content a A_280_, and fractions with protein were pooled and filtered with a 0.22 µm polyethersulfone syringe filter. Next, the pooled eluent was desalted to the equivalent of 20% (*i.e.,* ∼400 mM KCl) Heparin B buffer (25 mM HEPES-KOH pH 7.5, 2 M KCl, 10% glycerol, 1 mM DTT). Heparin A buffer (25 mM HEPES-KOH pH 7.5, 10% glycerol, 1 mM DTT) was used to dilute Heparin B buffer. The sample was loaded on to a 5 mL Heparin HP column (Cytiva) equilibrated in 20% Heparin B buffer. Unbound protein was removed with 20% Heparin B buffer. Bound protein was eluted in 15 column volumes with a gradient of 20% – 100% Heparin B buffer. Fractions were measured for protein content at A_280_, and those with protein were pooled and concentrated to 4 – 4.5 mL before filtration with a 0.22 µm polyethersulfone syringe filter. A SEC HiPrep 16/60 Sephacryl S300 column (Cytiva) equilibrated in SEC buffer (25 mM HEPES-KOH pH 7.5, 800 mM KCl, 10% glycerol, 5 mM DTT) was loaded with sample followed by one column volume of SEC buffer. Fractions in the size range of monomeric protein were measured for protein content at A_280_, and those with protein were pooled and concentrated to 8 mg/mL (50 µM) in SEC buffer using an Amicon centrifugal filter unit.

Aliquots were made at a volume no greater than 100 µL and were snap-frozen in liquid nitrogen and stored long-term at -80 °C. To make a working stock, the 100 µL 50 µM aliquot was rapidly thawed by directly transferring the tube from the -80 °C to a beaker of pre-warmed 37 °C water for <30 seconds. Once thawed, it was diluted to 10 µM by combining with 400 µL pre-chilled 4 °C storage diluent buffer (optimally 25 mM Tris-HCl pH 7.5, 200 mM KCl, 400 mM (NH_4_)_2_SO_4_, 0.2 mM TCEP, 50% glycerol; see main text for variations) and transferred to ice. The diluted enzyme stock was then carefully mixed by pipetting 50% of the volume ∼50 times on ice.

### OTTR cDNA library synthesis, cDNA purification, and cDNA size-selection

All oligonucleotides used were synthesized by Integrated DNA Technologies, with RNase-free high performance liquid chromatography purification included for oligonucleotides used in cDNA library synthesis. As described previously (Upton et al. 2021, Ferguson et al. 2023), to make Buffer 4B or Buffer 4B variants, primer-duplex DNA and RNA strands were first heated to denature structure at 70 °C for 5 minutes before annealing by decreasing the temperature to 4 °C with a -0.2 °C/second ramp. Annealed duplexes were made at 50 µM in water, diluted to 3.6 µM with additional water, and combined 1:1 with 7.2 µM 3′rC adapter template or 3′ddC adapter template also in water, resulting in a final Buffer 4B with 1.8 µM primer duplex and 3.6 µM adapter template.

All libraries in this study were synthesized using miRXplore Universal Reference Standard (Miltenyi Biotech, 130-094-407) as input RNA, or with no input template. OTTR v1 workflows and buffers are summarized in **Figure 1**. Input miRXplore miRNA was diluted to 9 or 10 µL in nuclease-free water. 2 µL of Buffer 1A, 1 µL of Buffer 1B or 1B2, and 1 µL of 10 µM working-stock BoMoC (either WT or mutant) or yPAP (Thermo Scientific, 74225Z25KU) were added sequentially. After 90 minutes at 30 °C (or other time, if indicated), 1 µL of Buffer 1C was added (or not, if indicated) and the reaction was either incubated for an additional 30 minutes (or not, if indicated) at 30 °C. 1 µL of Buffer 2 and 0.5 µL of rSAP (New England Biolabs, M0371S) pre-mixed with 0.5 µL of 50% glycerol were added. After 15 minutes at 37 °C, 1 µL of Buffer 3 was added, followed by 5 minutes at 65 °C, after which the reaction was placed on ice. 1 µL of Buffer 4A, 1 µL of Buffer 4B or Buffer 4B2, and 1 µL of the 10 µM working-stock BoMoC were added sequentially, followed by incubation for 30 minutes at 37 °C.

After OTTR, cDNA products were recovered largely as described previously (Ferguson et al. 2023). BoMoC was inactivated by incubation at 70 °C for 5 minutes, followed by addition of 1 µL RNase A (Sigma, R6513) and 1 µL of Thermostable RNase H (New England Biolabs, M0523S) before incubation at 55 °C for 15 minutes. Then 1 µL of Protease K (New England Biolabs, P8107S) and 30 µL of stop buffer (50 mM tris-HCl pH 8.0, 20 mM EDTA, 0.1% SDS) were added. After 15 minutes at 55 °C, the reaction was incubated at 95 °C for 5 minutes. Zymo Oligo Clean and Concentrator-5 columns were used for cDNA purification with elution in 6 µL nuclease-free water. 6 µL of 2X formamide loading dye (FLD; 95% formamide, 5 mM ethylenediaminetetraacetic acid (EDTA) pH 8.0, 0.05% (w/v) bromophenol blue, and 0.005% (w/v) xylene cyanol) was added. cDNA was denatured at 98 °C for 5 minutes and snap-cooled on ice prior to electrophoresis resolution on a 8% 19:1 acryl:bis-acryl 7 M urea 0.6X tris-borate-EDTA (TBE) dPAGE gel pre-run and run in 0.6X TBE buffer until the xylene cyanol dye-front was near the bottom of the gel. The gel was imaged to detect Cy5 and/or IRD800 using an Amersham Typhoon Trio (Cytiva) and the image was printed at 100% size of the gel to guide cDNA size-selection. Excised gel fragments were crushed against the sides of a 1.5 mL tube and cDNA was eluted into 400 – 500 µL of DNA elution buffer (300 mM NaCl, 10 mM Tris-HCl pH 8.0, and 1 mM EDTA) with a 70 °C incubation for 1 hour. cDNA was ethanol precipitated and resuspended in 30 µL nuclease-free water. cDNA was quantified by qPCR as described (Ferguson et al. 2023, McGlincy and Ingolia 2017) using iTaq™ Universal SYBR® Green Supermix (Bio-Rad, 1725120).

### Assays of 3′ labeling by dPAGE

In Figure 7A, 3′-labeling conditions which roughly matched the conditions during the 3′-labeling step in OTTR were used. Briefly, 6 µL of nuclease-free water, 2 µL of Buffer 1A, 1 µL of Buffer 1B2B (28 mM MnCl_2_, 28 mM (NH_4_)_2_SO_4_, and 10 mM sodium acetate pH 5.5), 1 µL of a 25 µM fluorescently labeled single-stranded DNA oligo (5IRD800/GTGACTGGAGTTCAGACGTGTGCTCTTCCGATCTNN), 1 µL of 10 µM working-stock W403AF753A BoMoC, and 3.5 µL of either 1.0 mM ddATP (MedChemExpress, HY-128036B), 1.0 mM dATP (ThermoFisher Scientific, 10216018), 1.0 mM dDAP-TP (TriLink Biotechnologies, N-2004), 1.0 mM ddATP-11-biotin (Revvity, NEL548001EA), 1.0 mM dATP-11-biotin (Revvity, NEL540001EA), 1.0 mM ddGTP-11-biotin (Revvity, NEL549001EA), or 1.0 mM dGTP-11-biotin (Revvity, NEL541001EA). For reactions with dATP-14-biotin, 0.75 µL of nuclease-free water and 8.75 µL of 0.4 mM dATP-14-biotin (ThermoFisher Scientific, 19524016) was used instead. After addition of the nucleotide triphosphate, the reaction was mixed by pipetting and incubated at 30 °C for 2 hours. Then, 1 µL of Protease K and 34.5 µL of stop buffer (50 mM Tris-HCl pH 8.0, 20 mM EDTA, 0.1% SDS) were added. After 15 minutes at 55 °C, the reaction was incubated at 95 °C for 5 minutes. Zymo Oligo Clean and Concentrator-5 columns were used for cDNA purification with elution in 6 µL nuclease-free water. 6 µL of 2X formamide loading dye (FLD, 95% formamide, 5 mM ethylenediaminetetraacetic acid (EDTA) pH 8.0, 0.05% (w/v) bromophenol blue, and 0.005% (w/v) xylene cyanol) was added. cDNA was denatured at 98 °C for 5 minutes and snap-cooled on ice prior to electrophoresis resolution on a pre-run 12% 19:1 acryl:bis-acryl 7 M urea 0.6X TBE dPAGE gel. The gel was resolved until the bromophenol blue dye-front was near the bottom of the gel. The gel was imaged to detect IRD800 using an Amersham Typhoon Trio (Cytiva).

### OTTR v2 cDNA library synthesis and streptavidin-based purification

OTTR v2 workflow and buffers are summarized in **Supplemental Figure 5**. Input miRXplore miRNA was diluted to 9 µL in nuclease-free water. 3′ labeling to extend the input template with a 3′ biotinylated ddATP was performed by adding 2 µL of Buffer 1A, 1 µL of Buffer 1B2A (1.0 mM ddATP-11-biotin; Revvity, NEL548001EA), 1 µL of Buffer 1B2B, and 1 µL of 10 µM working-stock BoMoC W403AF753A, sequentially. For ddRTP-11-biotin, 0.5 µL of Buffer 1B2A and 0.5 µL of ddGTP-11-biotin (Revvity, NEL549001EA) were used in place of 1 µL of Buffer 1B2A. After 120 minutes at 30 °C, or if specified 16 °C for 20 hours, 1 µL of Buffer 2 and 0.5 µL of rSAP pre-mixed with 0.5 µL of 50% glycerol were added. After 15 minutes at 37 °C, 1 µL of Buffer 3 was added. After 5 minutes at 65 °C, the reaction was placed on ice. Next, either 1 µL of Buffer 4A1, Buffer 4A4, or Buffer 4A was added (**Fig. 7C**). If Buffer 1B2A alone was used in the 3’-labeling step, 1 µL of Buffer 4B2 was added; for ddRTP-11-biotin, 1 µL of a buffer consisting of 3.6 µM c3t_ddCR5N adapter template, 0.9 µM of +T.v2.IR primer duplexed with RNA.v2.1 and 0.9 µM +1C.v2.Cy5 primer duplexed with RNA.v2.1 was used (**Table 2**, **Table 5**). Next, 1 µL of the 10 µM working-stock BoMoC was added and incubated for 30 minutes at 37 °C to complete cDNA synthesis.

Following cDNA synthesis, all cDNA purification steps were carried out at temperatures of 37 °C or below to ensure that the newly synthesized cDNA duplexes remained annealed. 1 µL of Protease K was added and incubated for 30 minutes at 37 °C. Double-stranded cDNA duplexes were purified from the 21 µL reaction by the addition of 63 µL of AMPure XP (Beckman Coulter, A63880) and incubation at room temperature for 10 minutes. The reaction was placed on a magnetic rack to immobilize the beads. After 5 minutes on the magnetic rack, the supernatant was removed and temporarily saved in case purification failed. While the tube remained on the magnetic rack, 200 µL of freshly-made 80% (v/v) ethanol was added to wash the beads. After 2 minutes the 80% ethanol was removed. This wash step was repeated a total of three times. Remaining 80% ethanol was carefully and completely removed from the beads, which were then left to air dry for 10 minutes. Completely removing all traces of ethanol was critical to avoid non-specific binding in a subsequent purification. 20 µL of nuclease-free water was added to the beads, the tube was removed from the magnet, and the beads were resuspended by pipetting. After 10 minutes of elution, the tube was returned to the magnet and the 20 µL eluent was removed and transferred to a new tube.

30 µL of Hydrophilic Streptavidin Magnetic Beads (New England Biolabs, S1421S) was added to a fresh tube on a magnetic tube rack and washed three times in Binding/Washing Solution (10 mM Tris-HCl pH 7.5, 1 mM EDTA, 1 M NaCl). After the final wash, 30 µL of Binding/Washing Solution and 20 µL AMPure XP eluent was added to the washed Hydrophilic Streptavidin Magnetic Beads. Streptavidin binding was performed by rotating the tube for 120 minutes at room temperature. After binding, the tube was returned to the magnetic rack, and after 5 minutes the supernatant containing unbound duplexes was transferred to a tube and temporarily saved to assess purification. A single wash was performed by adding 50 µL of Binding/Washing Solution. Elution was performed by adding 17 µL of nuclease-free water, 2 µL of 10X RNase H Buffer (New England Biolabs, M0297S), and 1 µL of RNase H (New England Biolabs, M0297S). RNase H hydrolysis of the biotinylated RNA template was carried out at 37 °C for 30 minutes. cDNA elution was completed by incubating the reaction at 95 °C for 5 minutes before transfer of the tube to a magnetic rack and removal of the eluent.

### OTTR library PCR amplification, purification, quantification, and pooling

For sequencing, 1 – 30 µL of cDNA was amplified by 6 to 12 PCR cycles using Illumina indexing primers and Q5 polymerase (New England Biolabs, M0491). For a typical miRXplore miRNA library from 500 ng input, 4 – 6 cycles were used. For libraries from 100 ng input, 10 cycles were used. For libraries from 20 pg input, 11 – 12 cycles were used. For libraries with less than 20 pg input, 12 – 14 cycles were used. PCR products were purified by DNA precipitation, Zymo DNA Clean and Concentrate columns, or AMPure XP (A63880, Beckman Coulter) following the recommendations of the manufacturer. DNA was resolved by 8% native PAGE and eluted as described (Ferguson et al. 2023). Eluted DNA libraries were quantified by qPCR using NEBNext® Library Quant DNA Standards (New England Biolabs, E7642S) and iTaq™ Universal SYBR® Green Supermix as described (Ferguson et al. 2023). Libraries were pooled based on these results. A maximum of 30 libraries were pooled at a time, with each library indexed by unique i5 and unique i7 sequences.

### Library sequencing and adapter trimming

All libraries were sequenced on an Illumina MiniSeq instrument using 75-(Illumina, FC-420-1001) or 150-(Illumina, FC-420-1002) cycle single-end high-output reagents. Libraries were denatured following Illumina recommendations. A final concentration of 1.0 – 1.2 pM denatured library in 500 µL of HT1 hybridization buffer (Illumina, 20015892) with 5% phiX (Illumina, FC-110-3001) was loaded to the sequencer. A sequential adapter trimming pipeline using cutadapt (v3.7) was used to trim FASTQ reads and retain crucial information about the reads (Ferguson et al. 2023). In detail, the adapter trimmed in the first step corresponds to a sequence encoded in the duplexed portion of the +1 primer duplex. In the second step, X nucleotide(s) were removed from the beginning of the read, with X=1 when no UMI (**Table 5**, c3t_NoUMI_6RNA) was used, X=4 when the 3′CNNN adapter template (**Table 5**, c3t_C3N_6RNA) was used, and X=7 when all other UMI-encoding adapter templates were used. In the third step, the final nucleotide that represents the primer-duplex +1 position was removed. In both the second and third steps, the sequences removed from the read were retained in the header of a FASTQ entry. In the final step, low-quality reads were trimmed and reads shorter than 15 nucleotides were removed.

cat untrimmed.R1.fastq | cutadapt -a
GATCGGAAGAGCACACGTCTGAACTCCAGTCAC - | cutadapt -u **X** –
rename=’{id} UMI={cut_prefix}’ - | cutadapt -u -1 –
rename=’{id}_{comment}_PD={cut_suffix}’ - | cutadapt -m 15 -q 10 - >
trimmed.fastq

### Sequence analysis

The miRXplore reference contains 962 synthetic equimolar miRNAs ranging from 16 – 28 nucleotides in length. The 943 subset of miRNAs in the 19 – 24 nucleotide range were used as a reference for read mapping. Across all libraries analyzed in this work, we calculated median read count for each miRNA; miRNAs with a median read count less than 50 across all libraries were not used to address differences in library depth, restricting the analysis to 904 miRNAs. CV was measured by dividing the standard deviation of the miRNA counts by the mean of the miRNA counts. For the analyses in Figures 2 and 3, during investigation of 5′ and 3′ precision of end-capture, we used a subset of 594 miRNAs based on their inability to misalign when the first and final three nucleotides were removed before mapping. End precision was measured by averaging the fraction of reads that included the first (for 5′ precision) or last (for 3′ precision) nucleotide for each miRNA (**Equation 1**).

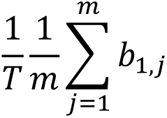

**Equation 1**: 5′ precision was determined by dividing the number of aligned reads to the first nucleotide (𝑏_1,𝑗_) of each miRNA (𝑗) by the total number of aligned reads (𝑇) and the number of miRNAs evaluated (𝑚 = 594).

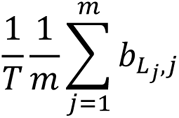

**Equation 2**: 3′ precision was determined by dividing the number of aligned reads to the final nucleotide (𝑏_𝐿𝑗,𝑗_, where 𝐿_𝑗_ is the length of miRNA 𝑗) of each miRNA by the total number of aligned reads (𝑇) and the number of miRNAs evaluated (𝑚 = 594).

In general, we allowed alignments as short as 15 nucleotides, but for the analyses in Figures 5 and 6, alignments shorter than 17 nucleotides were excluded for mapping confidence. All libraries were trimmed as described above before a sequential alignment pipeline using bowtie (v1.0.0) was performed. Trimmed reads were first aligned to a reference of OTTR oligonucleotide adapters. Unaligned reads were next aligned to the BoMoC expression-plasmid reference then the *E. coli* BL21 DE3 genome (NCBI AM946981.2). Filtering and sorting of alignments was performed by samtools (v1.7). miRXplore miRNA single-mapping alignments to both the reliable and unreliable miRNAs were converted to BED files by bedtools bamtobed (v2.25.0).

### Sanger sequencing of adapter-dimer cDNAs

Adapter-dimer-sized cDNAs were size-selected and eluted following 8% dPAGE as described above. Eluted cDNA was amplified by PCR using Illumina indexing primers as described above, but with 20 – 30 cycles and subsequent size-selection from a 2% agarose gel. Products were cloned into pUC19 vector.

### Assessment of *E. coli* nucleic acids co-purified with BoMoC

Size-selection used a cDNA size range of adapters plus 10 – 200 nucleotides. Eluted cDNAs for the 500 pg input libraries were amplified by 6 – 8 PCR cycles, while cDNAs for the 20 pg input libraries were amplified by 10 – 12 PCR cycles. Libraries were enriched by native PAGE size-selection of ∼160 – 350 bp duplexes. After sequencing and alignment as described above, the ratio of miRNA to *E. coli* and expression-plasmid reads was derived for all reads 18 nucleotides or longer. A line slope was computed from the ratio of reads mapped to miRXplore:contaminants and pg of miRXplore input.

### Statistical analysis

Comparisons of the fraction of precise 5′ or 3′ alignments were made by either unpaired one-sided or paired two-sided student’s *t*-tests performed only on miRNAs which had ≥30 alignments in both libraries being compared. For paired two-sided student’s *t-*test, the p-values were adjusted by Bonferroni correction based on the number of miRNAs analyzed and the results were represented as a mean difference in the fraction of precise 5′ or 3′ alignments. A result with an adjusted p-value less than 0.05 was defined as significant. Details and results from the paired two-sided student’s *t*-test were summarized in **Table 1** and **Table 5**.

### Dataset deposition

Illumina sequencing reads were deposited to the NCBI Sequence Read Archive under BioProject PRJNA1167688.

## Acknowledgements

H.E.U., L.F., S.C.P., and K.C. were supported by NIH grants R35 GM130315 and DP1 HL156819, as well as the Bakar Fellows Program (to K.C.) and NIH Grant T32 GM007232 (to L.F.). N.T.I. was supported by NIH grant R01 GM130996. We thank both the UC Berkeley DNA Sequencing Facility and the Vincent J. Coates Genomics Sequencing Laboratory QB3 Genomics, UC Berkeley, Berkeley, CA, RRID:SCR_022170, for sequencing support. We thank past and present members of the Collins and Ingolia labs of UC Berkeley for their support.

